# Comprehensive detection of structural variations in long and short reads dataset of French cattle

**DOI:** 10.1101/2025.05.22.654398

**Authors:** Maulana Mughitz Naji, Christophe Klopp, Thomas Faraut, Camille Eché, Arnaud Di Franco, Clément Birbes, Camille Marcuzzo, Amandine Suin, Carole Iampietro, Claire Kuchly, Caroline Vernette, Sébastien Fritz, Cécile Grohs, Christine Gaspin, Denis Milan, Cécile Donnadieu, Didier Boichard, Marie-Pierre Sanchez, Mekki Boussaha

## Abstract

Structural variants (SVs) correspond to different types of genomic variants larger than 50 bp. Many findings suggest the use of long rather than short reads to improve the accuracy of SV detection. Here, we present the results of an in-depth analysis for detection of SVs, mainly large insertions and deletions, in 14 French bovine breeds, based on whole-genome data comprising 176 long-read and 571 short-read samples, with 154 individuals having both long- and short-read data available. We first investigated possible biases on the performances of well-known SV detection tools, namely CUTESV, PBSV, and SNIFFLES, using long reads from different technologies, including PacBio HiFi, Oxford ONT, and PacBio CLR. We subsequently highlighted the abilities of tools for detecting SVs (DELLY, LUMPY, and MANTA) and for genotyping known SVs (GRAPHTYPER, SVTYPER, PARAGRAPH, and VG toolkit) using short-read data. We then show how the incremental composition of samples in the reference panel affected the SV genotyping for six validation individuals sequenced in short reads. We then searched for the optimal parameters and created the final SV reference panel consisting of 25,191 deletions and 30,118 insertions. Finally, we emphasized the landscape of the genotyped SVs segregating across 571 short-read individuals of 14 breeds.

## INTRODUCTION

Structural variations (SVs) are generally characterized as genomic alterations exceeding 50 base pairs (bp) and encompass various forms such as deletions, duplications, insertions, inversions, and translocations [1]. Due to their size, SVs are more likely to influence genetic variability than single nucleotide polymorphisms (SNPs) or small insertion-deletions (InDels) [1–3]. In contrast to SNPs and InDels, the development of SV detection pipelines presents greater challenges and requires further optimization. This complexity arises from factors such as the types of reads, the range of sizes, the methodologies employed, and the presence of multiple SVs at identical locations [1, 4].

Many studies have used short-read (SR) sequences to detect SVs, aligning them to a reference genome and using classical detections tools [5–8]. These SR-based tools often rely on alignment-related criterias, for instance on read-depth, split-reads, or abnormal distance between paired reads, from the alignment to detect SVs [9–11]. Recent studies indicated that this strategy is ineffective in accurately identifying structural variants (SVs). The low accuracy is attributed to the SR lengths and technical artifacts associated with SR alignment, especially in regions with repetitive sequences or high GC content [12, 13]. For example, half of deletions and three quarters of insertions cannot be detected using SR even with a combination of multiple variant callers [13]. Therefore, the use of long-read (LR) sequences is necessary as they allow for a more comprehensive SV detection [14, 15].

LR-based SV detection tools are mainly categorized into assembly or alignment based [15]. Briefly, the former assembles the reads *de novo* and then, compares this assembly to the reference genome to call SVs. In the alignment approach, LR sequences are mapped directly against the reference genome and SVs are detected according to different alignment signatures to the reference genome [4, 16]. It has been suggested that alignment-based methods generally have better genotyping accuracy than assembly-based methods, even with lower coverage [4, 15, 16]. Moreover, the alignment based allows, with no additional work, to detect heterozygosity whereas assembly based requires haplotype resolved assemblies to detect heterozygosity, which is not a trivial task [17, 18].

Several LR sequencing projects are underway, supported by the ever decreasing cost, creating an ideal situation where SVs will be discovered from long reads [4, 19, 20]. Currently, the main LR platforms are PacBio single molecule real-time (SMRT) and Oxford Nanopore Technologies (ONT) which produce reads length ranging from ten to several hundred kilobases [13]. In addition, PacBio protocols are categorized into continuous long reads (CLR; >30 kb and one polymerase pass) and high-fidelity (HiFi; 10-30 kb and multiple polymerases passes) based on the library insert size and number of polymerase passes over the fragment. This data type has emerged as the most prevalent in contemporary usage, owing to the benefits of both extended range and enhanced base accuracy (>99%) [13, 20]. Nonetheless, SR platforms account for a significant portion of genomic research, sequencing millions of genomes annually across various populations and species. This is primarily attributed to the reduced cost per base and the simplified library preparation process [4, 14]. Thus, SVs detection from SR might help us to better understand the causality of Mendelian and/or complex traits of interest. The challenge is how to make the best use of SR data given these discrepancies. In addition to classical SV callers from SR, there are recent tools available for genotyping known SVs using SR, such as GRAPHTYPER [21], PARAGRAPH [22], SVTYPER [23], and Variation Graph toolkit (VG) [24]. The different variants at a given locus are embedded in a graph, each subpath corresponding to a given allele. SRs are then re-aligned to this graph, and alignment evidences at and around the breakpoints are used to genotype the SVs [22, 25]. Using simulated data, previous studies have benchmarked the performance of various SR-based SV callers and genotypers with a presumably known truth-set of SVs [26, 27]. Nonetheless, the evaluation of SV detection using current datasets, especially in the context of livestock, remains quite restricted [28].

The reference bovine genome, which spans approximately 2.7 gigabases with thousands of recognised breeds, may contain a vast number of common and rare SVs that have yet to be discovered [19, 29]. Early work on the detection of 6,462 SVs in three major French breeds was carried out using SR whole genome sequence (WGS) data of 62 bulls [5]. Using SR data, another study detected approximately 68,000 SVs from 310 Holstein cattle of which approximately 10% overlap with the current BovineHD SNP array [30]. Another study reported association of three and twelve SVs toward several fertility traits in Holstein and Nordic Red Dairy cattle, respectively [31]. While another SV has an antagonistic effect on fertility and milk production traits [32]. Identified SVs in cattle were also found to be associated with gene expression, such as *SLC13A4 and TTC7B* in testis tissue [33]. In a similar manner, various SVs are notably linked to gene expression, which leads to phenotypic diversity associated with high altitude and coat color in domestic yak [34].

In light of these circumstances, the aims of the present study are to assess the effectiveness of various SV detection approaches and their potential implication in cattle population studies. This includes i) demonstrating the impact of different LR technologies and callers on SV detection; ii) highlighting the performance of different SV callers and genotypers using SR information; iii) exploring the effect of incrementing the number of reference samples and of applying different thresholds on the performance of the genotyping tools; and finally, iv) using the information of genotyped SV for population-based analysis in a larger SR dataset of French cattle.

To date, this study represents the most thorough examination of SVs, utilizing extensive LR and SR data derived from current cattle datasets. Consequently, our findings illustrate the potential practical uses of SV detection and genotyping within the bovine genome, and possibly in other species, to further investigate its relationship with the genetic variability of significant Mendelian and complex traits.

## RESULTS

### Evaluation of SV detected from different callers and LR data

We took advantage of a single Charolais heifer sequenced with three long-read (LR) technologies (see sup. Table1), with coverage up to 40x for HiFi, 52x for ONT and 42x for CLR [35]. We deemed these coverage levels adequate for a comprehensive analysis in light of the criteria established by earlier studies [18, 20] for assembly and variant discovery. After the alignment of reads from each technology to the ARS_UCS1.2 reference genome, SVs were detected using CUTESV [36], PBSV [37], and SNIFFLES [38], commonly-used software packages.

First, we characterized the SV detection results on HiFi data. After quality control, CUTESV, PBSV, and SNIFFLES detected 11,002, 11,575, and 13,567 deletions, respectively (sup. Fig 1). We conducted TRUVARI [39] benchmarking to identify intersections of SVs detected by the tools. For insertions, CUTESV, PBSV, and SNIFFLES detected 11,759, 10,258, and 13,913, respectively. Noteworthy, 10,000 deletions and 8,866 insertions, were commonly detected by the three tools (Fig 1a). We also observed that SNIFFLES identified 4,328 specific SVs, which was significantly higher than those detected by CUTESV (784) and PBSV (566).

**Figure 1:**
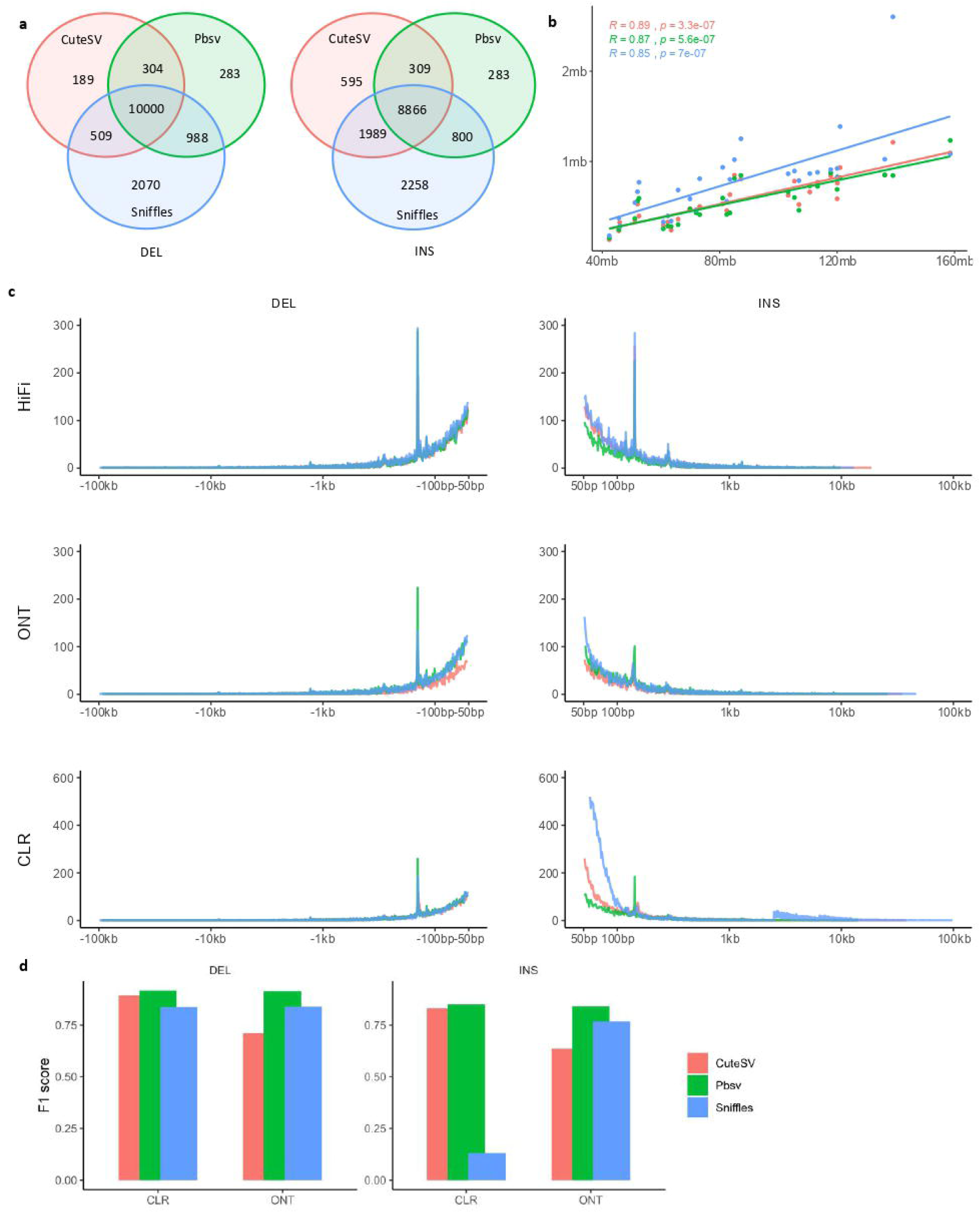
Properties of structural variants (SVs) discovered by various long-read technologies with different software on the genome of a single Charolais heifer. A) Venn diagram indicating the number of overlapping SVs called by CUTESV, PBSV and SNIFFLES software on the heifer’s PacBio-HiFi data. B) Plot of total number SVs discovered by the three callers against chromosome size. C) Frequency of discovered SVs according to their length (x-axis) for deletions (left) and insertions (right) using the three tools. The first, second, and third rows indicate the long-read sequencing of Pacbio-HiFi, Oxford-ONT, and PacBio-CLR, respectively. D) F1 score output from the truvari bench. The base set is the SVs called from HiFi and the comp set is the SVs called from ONT or CLR data using the CUTESV, PBSV and SNIFFLES callers.

The total numbers of SVs called by all three tools were highly correlated (R coefficient ≥ 0.85, Spearman’s correlation test) with chromosome size (Fig 1b). Heterozygous [0/1] and homozygous alternative [1/1] SVs were observed for this heifer called by the three tools. Although the fraction is relatively small, SNIFFLES also reported 99 insertions with uncalled genotype [./.] and 1,764 deletions and 1,827 insertions with homozygous reference [0/0] (sup. Fig 1, sup. table 2).

A symmetric distribution of SVs based on their length, between deletions and insertions, is expected [20, 35, 40]. For the HiFi data, the SV distribution followed this pattern between deletions and insertions across the three callers (Fig 1c). We observed a trend where the number of detected SVs declined as variant size increased. Clear peaks around 143 bp were detected for deletions (345, 287, and 294 for CUTESV, PBSV and SNIFFLES, respectively) and for insertions (256, 225, and 284 for CUTESV, PBSV, and SNIFFLES, respectively) (sup. Fig 2). For ONT data, the distribution pattern remained similar, but the peaks were lower and became less pronounced, especially for insertions, compared to HiFi data. For CLR data, the deletions length distribution mirrored that of HiFi and ONT data. However, the distribution pattern for insertions changed significantly, with an inflation of insertions around 50 to 100 bp, detected by CUTESV and SNIFFLES. In contrast, PBSV consistently detected the peak around 143 bp for both deletions and insertions across HiFi, ONT, and CLR technologies (Fig 1c, sup. Fig 2). Notably, this inflation of insertions was found not only in the Charolais heifer, but also in almost all individuals sequenced by PacBio CLR (sup. Fig 3-5).

To further evaluate the performance of CUTESV, PBSV and SNIFFLES across different LR data, we performed a benchmark between SVs called from ONT or CLR data (as the comp set) with the base set, SVs called from HiFi data. PBSV outperformed CUTESV and SNIFFLES with F1 scores above 0.91 for deletions and above 0.84 for insertions in both the ONT and CLR datasets (Fig 1d, sup. table 3). This indicates that PBSV calls showed the highest consistency in SV calling across HiFi, ONT and CLR data.

We further evaluated whether an ensemble calling, enriching the SVs discovered by PBSV with SVs called by CUTESV and/or SNIFFLES, could improve the benchmarking results. In this study, we utilized both the intersect and union approaches for the pairs of PBSV-CUTESV, PBSV-SNIFFLES, and a combination of three callers, applying them to CLR data as the comparison set in relation to the base set (see Methods). Our analysis revealed that none of the combinations achieved an F1 score that surpassed the default PBSV on CLR vs. PBSV on HiFi comparison (0.91 for deletions and 0.84 for insertions) (sup. table 4). Thus, we did not identify any further benefits in enriching SVs discovered by PBSV on CLR data with SV calls from other tools.

PBSV recommends using additional information from tandem repeat regions for increasing sensitivity and recall of SVs. We annotated tandem repeats and used this annotation during SVs calling in three HiFi samples from the Abondance (ABO), Tarentaise (TAR), and Vosgienne (VOS) breeds (sup. table 5). We compared the results to SVs previously discovered without annotation. The average F1 score was 0.98 corresponding to a relatively low number of SVs (∼300) that were different without tandem repeat annotation (sup. table 5). Consequently, considering the balance between the potential gains from SVs and the computational resources needed, we proceeded with the analysis using SVs identified by PBSV on LR data, excluding tandem repeat annotations.

At this stage, considering the benchmarking results for the Charolais heifer and the additional CLR individuals available in our dataset (sup. table 1), we used SVs detected by PBSV for the subsequent analysis in this study regardless of whether the sample was sequenced with HiFi, ONT or CLR.

### Evaluation of short reads SVs genotyping tools

We evaluated the feasibility of using SR data to recover the genotypes of identified SVs that were originally discovered through LR data. Performance evaluation metrics were derived from the F1 and genotype concordance scores produced by TRUVARI. We used both information to measure the ability of the tool to correctly detect SVs with correct genotype. The analysis included three main groups of tools based on read alignment and requirement of known SVs input information: 1. DELLY [10], LUMPY [11], and MANTA [9] which use evidence from the BAM file alignment to call SVs; 2. GRAPHTYPER [21], PARAGRAPH [22], and SVTYPER [23] which use the BAM file alignment and known SVs for local re-alignment nearby breakpoints to genotype SVs; 3. Variation Graph toolkit (VG) [24] which creates a genome-wide variation graph from SVs and aligns the entire reads to the graph for genotyping the SVs (Fig 2a).

**Figure 2:**
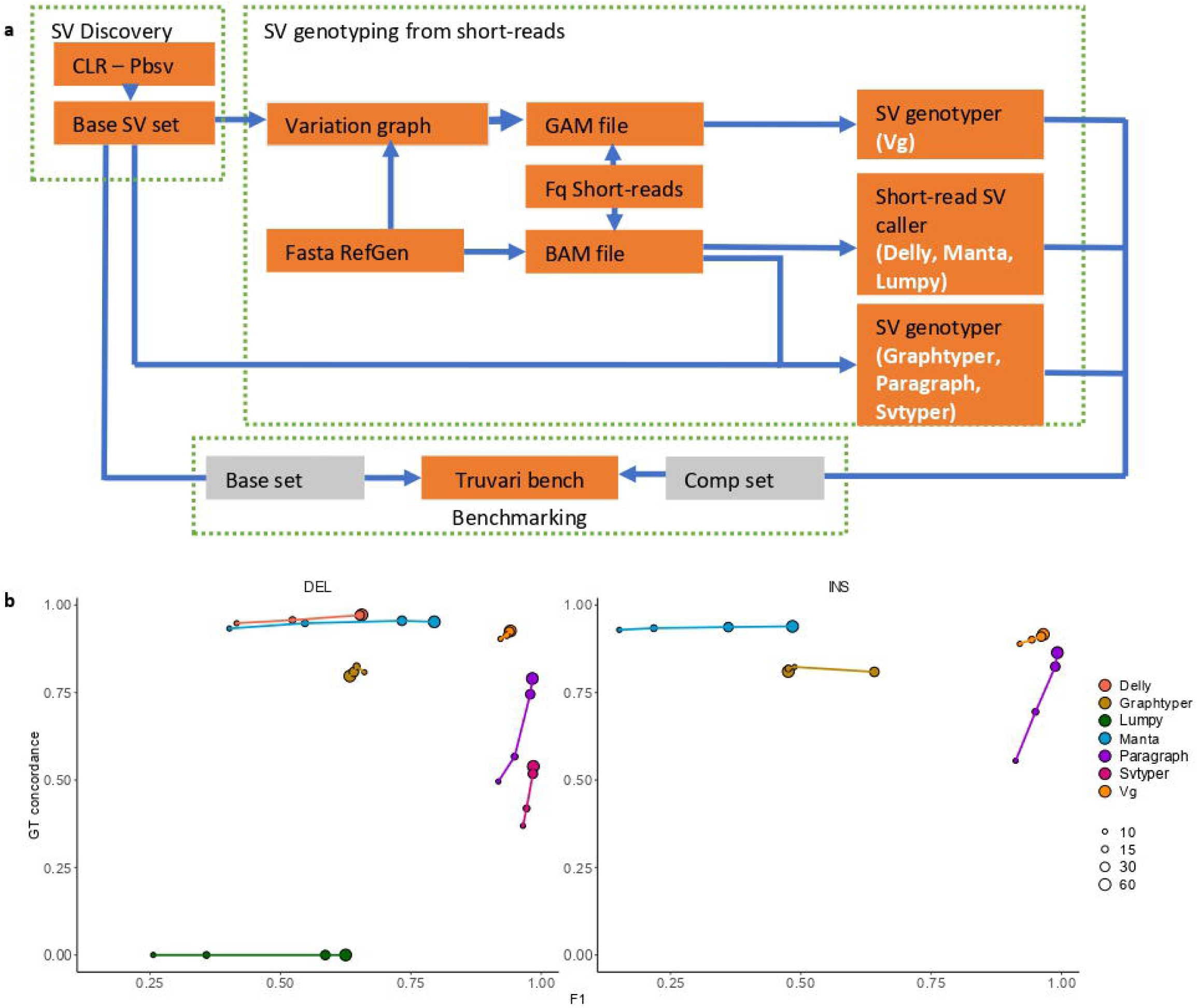
Structural variant (SV) genotyping of known SVs from short-reads. A) Workflow from benchmarking SV genotyping tools. B) Performance of SV genotyping tools on a single Charolais heifer for deletions (left) and insertions (right) using F1 indicator and genotype concordance scores at different levels of short-read coverage (circle size).

Again, we used the Charolais heifer, for which both LR and SR are available at varying coverage levels (see *Methods*). The F1 scores of the genotyping tool were higher when using SV information (Fig 2b). For example, on deletions, the F1 score of DELLY and LUMPY were both 0.63, comparable to GRAPHTYPER at the highest coverage of 60x. In contrast, MANTA’s score was 0.79, while genotyping tools using input SV information, such as VG, PARAGRAPH and SVTYPER, achieved F1 scores higher than 0.94. Note that LUMPY only reports SV breakpoints without genotype information. Only MANTA, GRAPHTYPER, SVTYPER and VG genotype insertions. Their F1 scores for insertions were lower compared to deletions, 0.48 for MANTA and 0.47 for GRAPHTYPER. Meanwhile, PARAGRAPH and VG slightly increased their F1 scores from 0.98 to 0.99 and 0.94 to 0.96, for deletions and insertions, respectively.

The performances of the genotyping tools were predominantly influenced by the coverage, with the notable exceptions of GRAPHTYPER and VG. For both deletions and insertions, the coverage affected the F1 scores of DELLY, LUMPY and MANTA. The genotype concordance of paragraph and SVTYPER were also affected by the coverage. Furthermore, considering that the SR alignments are carried out independently for each SV genotyping tool, the resource requirement is better optimized for VG because it only used one third of the time (92 cpu-hours) compared to the other genotyping tools (330 cpu-hours) for the SR alignment with 60x coverage (sup. table 6).

We expanded the evaluation of SV genotyping tools by conducting an independent analysis of 154 samples with both LR and SR available (Table 1, sup. table 1). The comp sets included SVs that were genotyped using various tools with SR data, whereas the base set consisted of SVs called by pbsv from CLR data (Fig 2a). After removing the outliers, 148 samples were retained in the analysis. In this group of individuals, a greater number of deletions were identified compared to insertions (Fig 3a). The number of SVs differed significantly, ranging from 15,472 in HOL19 to 23,086 in AUB3.

**Figure 3:**
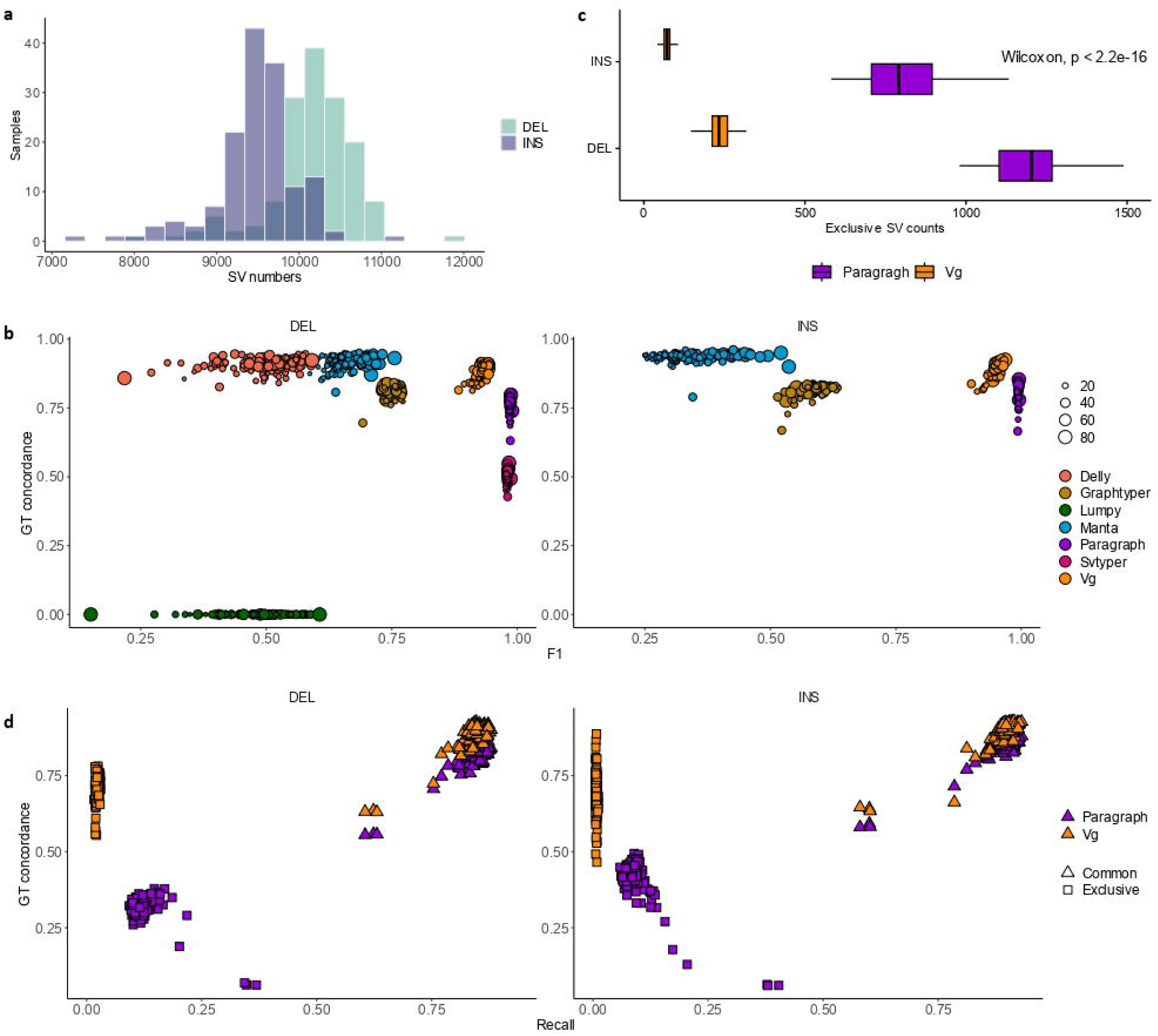
Structural variant (SV) genotyping from short-reads in 148 individuals with long and short-reads available. A) Histogram indicating the number of SVs in the base set called by PBSV on CLR data. B) Performance of SV genotyping tools on deletions (left) and insertions (right) based on F1 and genotype concordance scores. C) Boxplot with Wilcoxon rank test on the number of SVs genotyped exclusively by PARAGRAPH and VG. D) Recall and genotype concordance scores of commonly (intersect) and exclusively genotyped SVs by PARAGRAPH and VG against the base set of the respective individuals.

**Table 1.** Summary of long-read (LR) and short-read (SR) dataset used in the study.

| Breed | Abbreviation | LR samples | LR and SR available | Target SR samples |
| --- | --- | --- | --- | --- |
| Abondance | ABO | 12 | 11 | 13 |
| Aubrac | AUB | 12 | 11 | 40 |
| Blonde d'Aquitaine | BAQ | 11 | 10 | 31 |
| Brown Swiss | BSW | 5 | 5 | 9 |
| Charolaise | CHA | 17 | 10 | 60 |
| Holstein | HOL | 31 | 25 | 147 |
| Hybrid* | - | 2 | - | - |
| Limousine | LIM | 10 | 10 | 88 |
| Montbéliarde | MON | 26 | 26 | 91 |
| Normande | NMD | 28 | 26 | 64 |
| Parthenaise | PAR | 3 | 3 | 4 |
| Rouge Flamande | RDC | 3 | 3 | 5 |
| Simmentale | SIM | 5 | 5 | 5 |
| Tarentaise | TAR | 6 | 5 | 10 |
| Vosgienne | VOS | 5 | 4 | 4 |

\*Two hybrid samples were crossed of HOL with NMD and Yak with MON, respectively.

The performance of the genotyping tools was consistent with observations made on the individual Charolaise heifer (Fig 3b). Tools were grouped similarly based on both F1 and genotype concordance scores, reflecting coherent clustering patterns. For F1 scores, PARAGRAPH, SVTYPER, and VG demonstrated significantly superior performance in genotyping deletions compared to the other tools (Fig 3b, sup. Fig 6). For insertions, PARAGRAPH and VG showed significantly better performance compared to the other tools (sup. Fig 7).

We conducted an additional comparison of SVs identified by both PARAGRAPH and VG, leading to the classification of SVs into four distinct sets for all individuals: intersect SVs genotyped by VG, intersect SVs genotyped by PARAGRAPH, exclusive SVs genotyped by VG, and exclusive SVs genotyped by PARAGRAPH. Each SV set was subsequently compared to the base set of the respective individual. It is noteworthy that there were as many as five times more SVs uniquely genotyped by PARAGRAPH compared to VG (Fig 3c). When categorizing the genotyped SVs into groups, the precision and recall rates of each group became less informative due to their significantly reduced numbers in comparison to the base set. Thus, the primary objective of this comparison was to determine the accurate assignment of heterozygous [0/1] or homozygous alternative [1/1] genotypes as established in the base set. Our analysis revealed that VG exhibited superior genotype concordance compared to PARAGRAPH for both intersecting and exclusive SVs, encompassing deletions and insertions (Fig 3d). Across 148 samples, the average genotype concordances scores were 0.78 for PARAGRAPH and 0.89 for VG.

In light of the independent benchmarking results concerning the Charolais heifer and the other 148 individuals, we have determined that VG is the most performing genotyping tool for recovering SVs from SR and will be utilized in the forthcoming analyses.

### Extending the number and diversity of reference SV panel

In the previous section, we assessed the performance of SV genotyping tools on a sample-by-sample basis, with VG achieving the best results. In this section, we investigated the influence of incorporating samples from various breeds into the variation graph on the performance of SV genotyping by VG.

We selected a subset of 151 samples, for which both LR and SR were available (Table 1, Sup. table 1). This subset was subdivided into 145 reference and 6 validation samples, comprising two Holstein (HOL), two Montbéliarde (MON), and two Normande (NMD) individuals. The workflow (sup. Fig 8) began with merging the base sets of reference samples using JASMINE [41], followed by the construction of the variation graph (see *Methods*). Validation individuals were then genotyped using SR mapped against the variation graph, and the resulting genotyped SVs were compared to the base set of each respective validation individual. The number of reference individuals used to construct the SV panels was gradually increased from 23, 46, 70 to 145, corresponding to 1, 2, 3, and 14 breeds, respectively. As the number of reference samples increased, the total number of SVs with distinct alleles and their corresponding bubbles in the graph also increased (sup. table 7). Across the four panels, 94.4% of the bubbles created were biallelic, while the remaining bubbles were multi-allelic resulting from SVs with identical breakpoints but multiple alternative alleles. These multi-allelic bubbles were already present in the single-breed panel but became increasingly abundant as additional reference breeds were added (sup. table 8). For example, a bubble indicating an insertion at chr 25: 39,613,777 initially appeared as biallelic, with one alternative insertion allele in panel one (HOL only panel). This remained unchanged in panel two (HOL + MON panel). However, in panel three (HOL + MON + NMD panel), the bubble became more complex by acquiring an additional allele common to NMD samples. By panel 14, which included 14 breeds as references, this bubble had three alternative alleles (Fig 4a). Overall, the number of bubbles almost doubled from panel one to panel 14, reflecting the increased diversity introduced by the broader breed representation (Fig 4b, sup. table 7).

**Figure 4:**
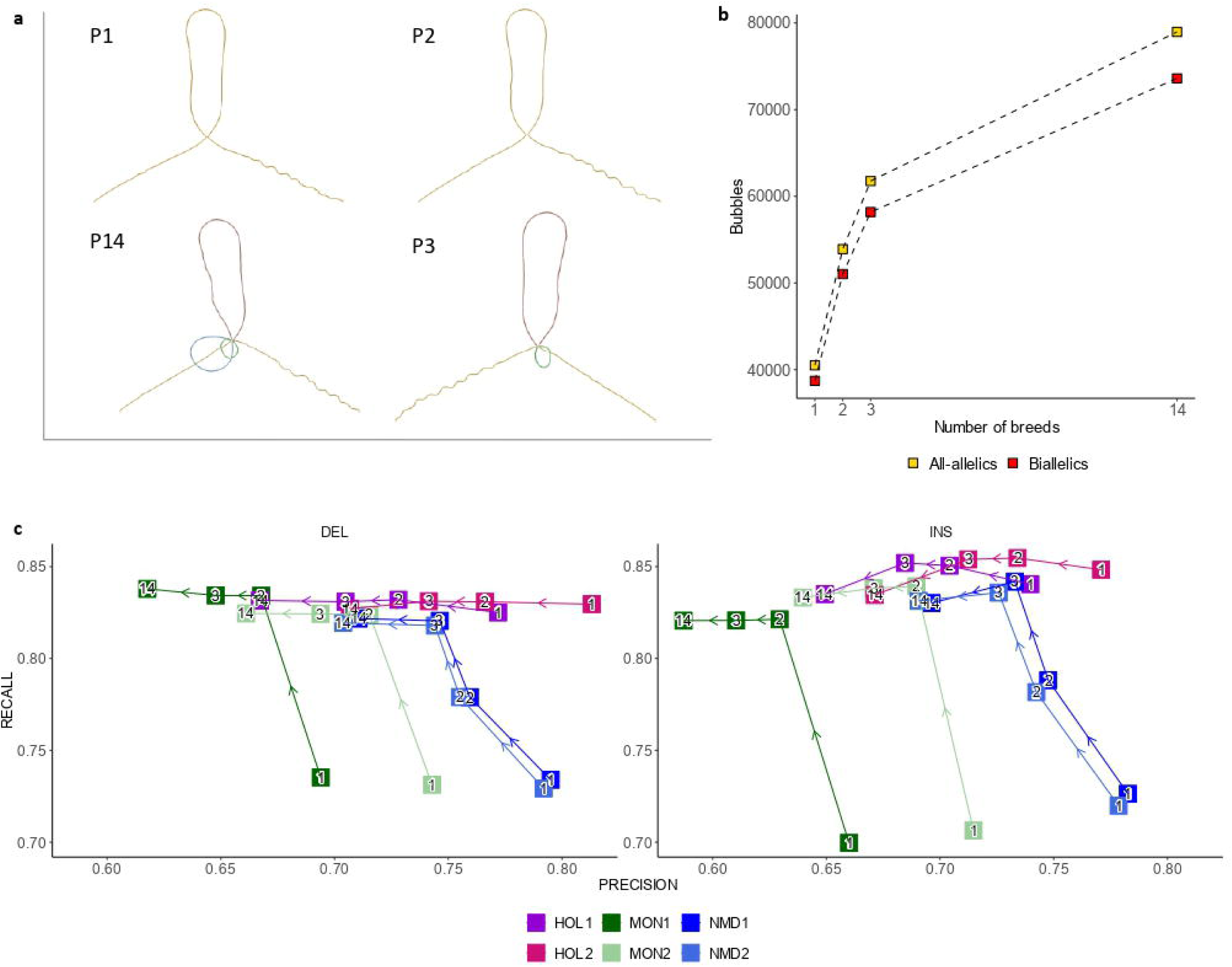
Increasing the number of reference samples in the structural variant (SV) panels affects the genotyping performance. A) The variation graph bubble indicates the insertion node at position 25: 39613777 created from *(in clockwise)*: P1 includes 23 HOL samples; P2 includes 46 HOL and MON samples; P3 includes 70 HOL, MON, and NMD samples; P14 includes 145 samples from 14 breeds. B) Increasing number of bubbles in the variation graph as the number of reference samples increases. C) Benchmarking of genotyped SV of 6 validation individuals from short-reads on different variation graphs. Left and right columns represent deletions and insertions, respectively. The values in the graph dots indicate the number of breeds on which the variation graph was created. Breed abbreviations are: HOL (Holstein), MON (Montbéliarde), NMD (Normande). The other 11 breeds are Abondance, Aubrac, Blonde d’Aquitance, Brown Swiss, Charolais, Limousine, Rouge Flamande, Parthenaise, Simmental, Tarantaise, and Vosgienne.

The precision and recall rates of the validation samples were shifted depending on the reference panels used to construct the variation graphs (Fig 4c). In panel one, the two HOL samples achieved the highest recall rates, approximately 0.82 for deletions and 0.84 for insertions. The inclusion of 23 MON into the reference for the creation of panel two improved the recall rates of two MON samples (Fig 4c, sup. table 8). For example, MON1’s recall rate for deletions improved from 0.73 in panel one to about 0.83 in panel two. Similarly, the NMD sample’s recall rates also improved from around 0.73 to 0.78 for deletions and from around 0.72 to 0.78 for insertions in panel two. When NMD samples were then added into the reference to form panel three, their recall rate further improved compared to when this breed was excluded. Specifically, the recall rates of two NMD samples reached 0.83 for the deletions and 0.84 for insertions in panel three. As more breeds were added into the reference panel, recall rates slightly increased and became more consistent across validation samples. For example, recall rates for the HOL, MON, and NMD validation samples on panel 14 were around 0.83 for both deletions and insertions.

On the other hand, accuracy rates tended to decline as more breeds were added to the reference panel. This pattern was observed uniformly among all validation individuals, showing an average decrease of 0.08 in precision rate from panel one to panel 14. For example, HOL1 had a precision rate of 0.77 on panel one and only 0.67 on panel 14. As a side note, no clear trend was observed in genotype concordance rates as the reference panels evolved (sup. Fig 9).

We deduced that incorporating individuals from particular breeds into the reference panel facilitates the genotyping of breed-specific SVs, thereby improving recall rate. Nevertheless, as the number of breeds in the panel increases, VG was compelled to genotype SVs that were absent from the validation samples, leading to a reduction in precision.

### Optimum parameters for building an SV reference panel of 14 breeds

Here, we used the same set of reference and validation samples as described in the previous section. We merged SVs from 145 samples, resulting in 107,274 and 126,008 SVs with distinct alleles using the default and adjusted parameters of JASMINE, respectively (see Methods). Subsequently we implemented eight VARCALLS which represented the number of variant calls that supported the merged SVs, of 2, 3, 5, 7, 13, 20, 27, and 36 as filters to the two merged SV sets. This process resulted in the creation of 16 reference panels. These SV panels served as the basis of constructing variation graphs on which SR of validation samples were then aligned.

F1 scores increased linearly with the application of more stringent VARCALLS thresholds (sup. Fig 10). Higher VARCALLS thresholds reduced the number of SVs to be genotyped, thereby increasing the precision rate (sup. Table 10). In contrast to precision, recall rates across validation individuals peaked between thresholds of five and seven, when deletions and insertions were considered together (Fig. 5a). By separating deletions and insertions, the optimal VARCALLS threshold was determined to be between seven and thirteen (sup. Fig 11). Therefore, we selected a VARCALLS threshold of seven as the optimal criteria for filtering. As a result, only SVs detected in at least seven samples were retained in the reference panel. At the same VARCALLS threshold, employing the adjusted merging parameter led to an increase in the number of SVs in the reference panel when compared to the default parameters (sup. table 9). However, the difference between the numbers of input SVs and bubbles were significantly higher with the adjusted merging parameters of JASMINE in the comparison to the default (Fig. 5b). Furthermore, within a VARCALLS of seven, the number of biallelic bubbles in the variation graph of the default merging was higher (52,053) than with the adjusted parameters (51,576) (sup. table 9).

**Figure 5:**
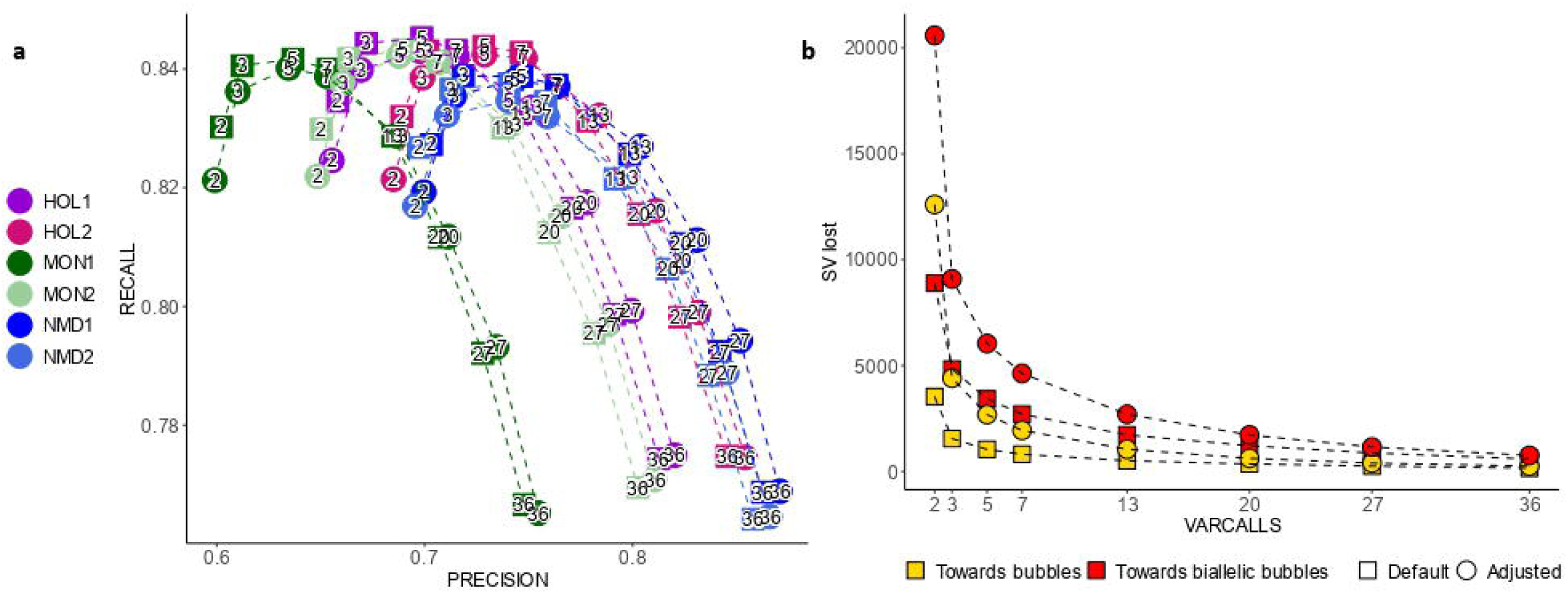
Finding optimal parameters for creating a reference panel of structural variants (SVs) from 14 breeds. A) Benchmarking of genotyped SVs from 6 validation individuals on the reference panel created from 145 samples from 14 breeds. JASMINE SVs merging was applied with default and adjusted parameters combined with different VARCALLS thresholds of 2, 3, 5, 7, 13, 20, 27, 36 as indicated by the points in the graph. Square and round dots indicate default and adjusted JASMINE merging parameters. B) Number of input SVs not translated into bubbles or biallelic bubbles in the variation graphs across VARCALLS threshold.

Taking this into consideration, we sought to identify the optimal parameters that would achieve an optimal balance between the recall rate and the total number of SVs within the fourteen-breed’s panel. Based on our evaluation of the validation samples, we considered that merging with the default JASMINE parameters and applying VARCALLS threshold of seven provided a satisfactory balance for creating the all-breed panel when merging from the base set of all LR samples.

### Genotyping SVs on a large cohort of individuals sequenced with short reads

Considering the recommendations from the previous section, we merged the base sets of 176 samples from 14 breeds (Table 1) using the default JASMINE parameters, resulting in 159,792 structural variations (SVs). After applying the VARCALLS threshold of seven, we retained 58,427 SVs. This filtered set was then used as input to construct the variation graph, which contained 55,309 biallelic bubbles representing 25,191 deletions and 30,118 insertions (sup. table 10). In total, these SVs provided alternative of 26,538,017 bp through deletions and 25,495,462 bp through insertions to the bovine reference genome.

The length distribution of SVs in the variation graph was similar for deletions and insertions, indicating a high-quality SV panel (Fig. 6a). However, we observed that deletions tended to be longer than insertions. Despite this, the graph contained a higher number of insertions than deletions (sup. table 10). Based on this graph, we then performed genotyping of SVs for 571 SR samples (Table 1). Across individuals, the average number of genotypes were 377 uncalled [./.], 32,645 homozygous-reference [0/0], 14,535 heterozygous [0/1], and 7,750 homozygous-alternative [1/1] (sup. table 11). Accordingly, the uncalled genotypes were masked as homozygous-reference.

**Figure 6:**
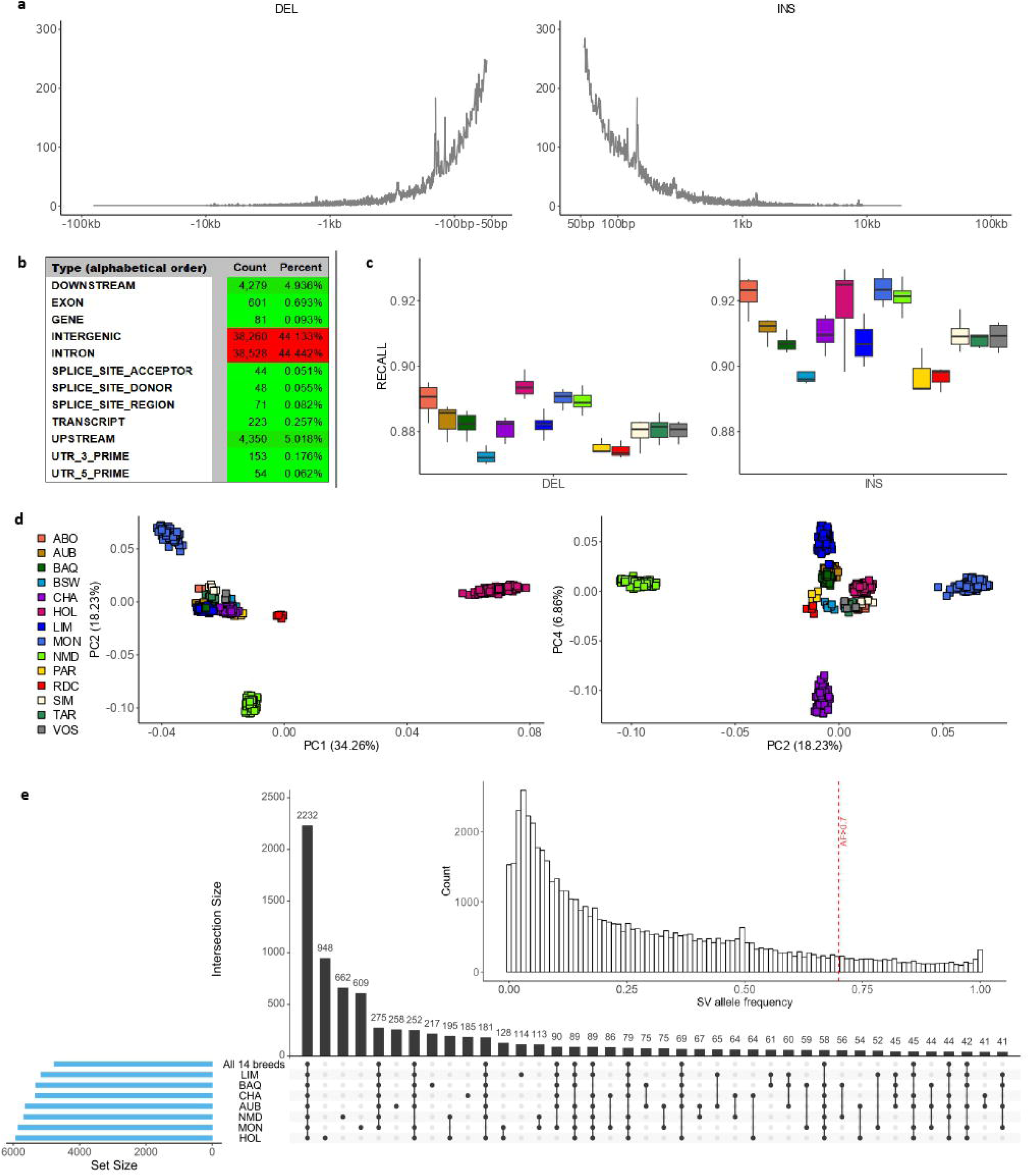
Genotyping of structural variations (SVs) in 571 short-read samples using a variation graph constructed from 176 long-read samples. A) Length distribution of genotyped SVs from the variation graph, showing deletions (left) and insertions (right). B) Annotation of genotyped SVs based on their genomic region type. C) Recall rates of genotyped SVs for 148 samples with both short-read and long-read data. D) Principal component analysis (PCA) of genotyped SVs across 571 short-read samples. E) Upset plot of major SVs (allele frequency > 70%) across all 14 breeds and 6 breeds with more than 30 short-read samples. The inset displays SV allele frequencies across 571 samples, with a vertical line indicating the threshold for major SVs. Breed abbreviations: AUB (Aubrac), BAQ (Blonde d’Aquitaine), CHA (Charolais), HOL (Holstein), LIM (Limousine), MON (Montbéliarde), NMD (Normande).

Most of genotyped SVs were located in intergenic (44.13%) and intronic (44.42%) regions, while 5.01% were upstream and 4.97% downstream of known genes (Fig. 6d). Only a small fraction (0.69%) was found within exons. Using snpEff [42], 86,692 effects were associated with these SVs, with 98.86% classified as having a modifier effect and less than 1% categorized as high impact (sup. Fig 12). These high impact SVs comprised of 370 deletions and 62 insertions overlapping with 368 protein coding genes and 55 RNA-related regions. On average, a genotyped SV was observed approximately every 47 kb.

On 148 samples with both LR and SR available, we benchmarked their genotyped SVs against their base sets from LR. For each breed, the average recall rates were higher for insertions compared to deletions (Fig. 6b). For instance, the difference between deletion and insertion recall rates were around 0.03 across ABO, HOL, MON, and NMD. No clear difference in recall rates was observed between the breeds due to different size of reference or genotyped samples. For instance, the average recall rate of insertions on ABO was 0.922 while HOL, MON, and NMD had an average recall rate of 0.923, 0.920, and 0.921, respectively. Average recall rates of breeds with reference samples below seven, corresponding to VARCALLS filter of seven, were among the lowest, such as the BSW (n=5), PAR (n=3), and RDC (n=3). Yet, recall rates of SIM (n=5), TAR (n=6), and VOS (n=5) were comparable to breeds with higher number of reference samples, for instances to LIM (n=10) and CHA (n=18). This suggests that most of breed-specific SVs belonging to BSW, PAR, and RDC were removed during the filtration step. In contrary, SVs found within SIM, TAR, and VOS were well-represented from the merging of 176 reference samples with the applied filter criteria.

Principal component analysis (PCA) of the genotyped SVs revealed a distinct differentiation among HOL, MON, and NMD when considering the first two components (Fig. 6c). The first component explaining 34.26% of the variance mainly separated HOL from the other French breeds. The second component, accounting for 18.23% of the variance, further separated MON and NMD at opposite extremes, while the remaining breeds closely clustered. Additionally, the fourth principal component, explaining 6.86% of the variance, further differentiated CHA, MON, NMD, and LIM. This result was consistent with the PCA performed on the SVs detected from 176 LR samples used as the reference for constructing the variation graph (sup. Fig 13).

Across 571 samples, the majority of SVs were found to be rare, with their frequencies primary clustered around 4-5% (inset of Fig. 6e). The frequency distribution of genotyped SVs revealed two distinct extremes. At one end, there were 350 deletions and 300 insertions that were homozygous-reference in all samples, indicating that no SV alleles were genotyped. Out of these SVs, seven were derived from samples with only LR sequences, while the remaining SVs were initially detected in the base set of 148 samples that have both LR and SR data (sup. Fig 14). Of the 650 SVs in this category, 341 (52.46%) overlapped with known repeat regions in the bovine reference genome, including LINE (36.60%), SINE (35.09%), LTR (11.12%), simple repeat (7.48%), and DNA transposons (5.05%). The GC contents of the reference (for deletions) or alternative (for insertions) sequences were significantly higher with a p-value < 2.2e-16 (unpaired wilcoxon-rank test), compared to all 55,309 genotyped SVs.

On the other hand, there were 106 deletions and 75 insertions that were homozygous-alternative across all samples. Out of these, 141 (46.17%) were found to overlap with the repeat regions, specifically LINE (46.17%), SINE (18.89%), LTR (16.26%), simple repeats (10.76%), and satellites (3.34%). Of note, 34,298 of 55,309 genotyped SVs overlapped with repeat regions, with proportion as follows: LINE (45.12%), SINE (28.21%), simple repeats (10.37%), LTR (2.95%), DNA transposons (1.39%), and satellites (1.39%). Here, GC contents of fixed-alternative SVs had the same distribution as all 55,309 genotyped SVs.

We excluded the two extreme SV categories and applied an arbitrarily allele frequency (AF) threshold higher than 0.7 to identify major SV alleles. Across 14 breeds, 2,232 SVs (4.03 % of total genotyped SVs) were classified as major alleles. HOL, MON, and NMD had the highest numbers of breed-specific major SVs, with 948, 609, and 662, respectively (Fig. 6e), exceeding those found in other breeds such as AUB (258), BAQ (217), CHA (185), and LIM (114). Some SVs were major in most breeds but absent in specific ones, including 275 SVs missing in HOL, 252 in NMD, and 181 in MON.

Similarly, 6,754 SVs (12.21 % of total genotyped SVs) were consistently rare (AF < 0.10) across all 14 breeds. HOL, MON, and NMD, had specific 1,505, 878, and 1,081 SVs, respectively (sup. Fig 15). Again, for the rare SVs, these three breeds surpassed those observed in AUB (406), BAQ (416), CHA (360), and LIM (209).

## DISCUSSION

A total of 176 samples were sequenced using LR technologies, either PacBio HiFi, ONT, or PacBio CLR, with the predominant technology being CLR. Therefore, we needed to identify the most reliable tool among CUTESV, PBSV, and SNIFFLES for consistently detecting deletions and insertions across all three sequencing platforms. Analysing a single heifer sequenced with all three technologies, we observed bias particularly for insertion detection in PacBio CLR data by SNIFFLES and CUTESV. This result was not unexpected, as the developers recommend using SNIFFLES primarily for ONT or HiFi data [38]. CLR, with its longer raw reads, is known to have higher error rates, mainly due to false insertions [4]. As a result, SNIFFLES detected more insertions in our CLR dataset.

For CUTESV, the F1 score derived from our heifer CLR dataset, which has approximately 42x coverage, was slightly lower than that reported for CLR of HG002 with 69x coverage, *i.e.,* 0.89 for deletions and 0.83 for insertions, instead of 0.90 [36]. In our ONT data, PBSV slightly outperformed CUTESV and SNIFFLES. This contrasts with the results observed in human NA24385 ONT data, where CUTESV (0.90) and SNIFFLES (0.87) outperformed PBSV (0.82) [43].

This finding supports the idea that tool performance is affected by various factors such as coverage, sequencing type, SV types, and the dataset or species being studied. In our study, PBSV demonstrated consistent performance for both deletions and insertions from HiFi, ONT and CLR reads.

We evaluated the ‘classical’ SR-based SV detection tools of DELLY, LUMPY, and MANTA against SV genotyping tools that also rely on SR data but incorporate a known SVs information from LR sequencing. Across deletions and insertions, the average F1 scores for these classical tools, based on an independent assessment of 148 samples, were approximately 0.50, significantly lower than genotyping tools using known SV information from LR, such as GRAPHTYPER (0.67), PARAGRAPH (0.99), SVTYPER (0.98), and VG (0.94).

Our results also suggested VG and PARAGRAPH as the most performing tools for SV genotyping, as evidenced by their F1 scores and capability to genotype both deletions and insertions, in agreement with other findings [25]. However, our average F1 scores were higher than those reported in 3 individuals of the Human Genome Structural Variation Consortium (HGSVC) with 0.72 for PARAGRAPH and 0.83 for VG [25]. We further demonstrated that VG outperformed PARAGRAPH in the genotyped concordance rate independently across the 148 samples. During our analysis, we also noticed that PANGENIE [40] is able to genotype SV from short reads. However, as its algorithm is based on assembly creation from long-reads, we excluded it from our analysis, which focused mainly on alignment-based methods.

We incrementally added reference samples from available breeds to create SV reference panels and subsequently re-genotyped these variants using SR from the 6 set-aside validation samples. In a human study, a leave-one-out approach was used also by generating reference variants from 10 samples while using one individual as a validation subject [40]. We illustrated the impact of reference size and the number of breeds on the performance of SV genotyping using VG. For example, the incorporation of individuals from a particular breed, (analogous to the concept of origin in humans) enhances the diversity of the reference panel and thus increases SV recall rate for validation samples from the same breed, even if they are not directly included as the reference. This improvement occurs because SVs that exhibit significant segregation within the breed are effectively represented by other individuals of that breed.

To determine the optimal parameters for constructing the reference panel encompassing all 14 breeds, we primarily focused on the recall rate. This metric indicates the efficiency with which SVs can be recovered in six validation samples that were excluded from the panel used to construct the variation graph. We did not prioritize the precision rate because all SVs in the graph were genotyped, and for SV sites without evidence of an alternative allele, a homozygous-reference genotype was assigned using VG with the *-Aa* parameter. Consequently, the substantial number of SVs in the graph resulted in a decreased precision rate, though the genotyping accuracy for SVs with alternative alleles remained unaffected. Therefore, we prioritized recall rate based on the six validation samples.

We chose a simple method using JASMINE to combine SVs across multiple samples, although other tools, such as BCFtools [44], SURVIVOR [45], and TRUVARI [39] could be used as well. Due to the extensive range of parameter combinations for merging SVs from reference samples using JASMINE, we restricted our analysis to two configurations: the default and adjusted parameters (see Methods). These were combined with eight VARCALLS thresholds to identify the optimal parameters, ultimately leading to the selection of JASMINE’s default merging parameters and a VARCALLS threshold of seven for constructing the final 14-breed panel.

We initially retained only biallelic variants from SV discovery using pbsv on LR from individual samples, yet the merging process eventually introduced multiallelic variants and, consequently, multiallelic bubbles in the generated variation graph. These multiallelic bubbles were observed in all reference panels, from the single-breed to the 14-breeds panel. They accounted for up to 3.5% of the 53,948 SV in our final graph of 14 breeds. In contrast, the human pangenome graph shows that multiallelic bubbles account for up to 89% of the 413.809 SVs [20]. The discrepancies in both SV counts and multiallelic bubbles can be attributed to the different sample sets used: their study included 47 different human assemblies, whereas we used 14 different cattle breeds, albeit with a larger sample size of 176.

The application of variation graph on 571 SR samples enabled genotyping insertions, even with higher recall rates than deletions across 148 samples, which was difficult to detect using the classical approach from SR data [46]. Thus, we expected these genotyped insertions, together with deletions, have potential implication to be discovered as many studies relied solely on the deletions for their association studies [31, 32, 47].

We observed two unexpected patterns in the genotype frequencies of SVs across all 571 samples. First, 650 SVs were consistently homozygous for the reference allele. These SVs exhibit relatively higher GC content than other genotyped SVs, which may have impaired short-read alignment to the variation graph, resulting in insufficient evidence to call alternative alleles in any sample. Second, several SVs showed a fixed allele frequency (AF) of one across all samples, suggesting that these variants may be specific to Dominette, the Hereford cow used to generate the ARS_UCD1.2 bovine reference genome.

These two extreme cases, which represent 1.5% of all genotyped SVs, cannot be solely attributed to repetitive, hard-to-genotype regions. In fact, 62% of the SVs in the final panel also overlap with repeat regions but were still successfully genotyped using short-read data in the bovine reference genome.

Overall, we successfully genotyped most SVs across individuals, achieving relatively high recall rates for both deletions and insertions. However, as previously noted, SV genotyping from SR data using variation graphs is not without limitations. LR sequencing remains essential for accurate SV detection—not only because some genotypes cannot be fully resolved using variation graphs alone, but also because SVs identified from LR data are required to construct the graph in the first place. Our findings thus support the notion that LR sequencing is indispensable for effective SV detection [14, 15]. Still, due to its higher cost and limited availability, the variation graph approach offers a practical and more scalable alternative to directly calling SVs from SR alignments.

Our study presents a systematic strategy for integrating LR and SR sequencing data to comprehensively detect structural variants (SVs) across diverse cattle breeds. The reference panel, built from SVs identified in 176 samples, and the resulting SV panel, genotyped across 571 samples from 14 breeds, together provide a valuable resource for population and association studies aimed at identifying genetic variants linked to important phenotypic traits.

## MATERIAL AND METHODS

### Detection of structural variations from long-reads

In this study, we used whole genome long read (LR) data available for 176 animals from 14 French dairy and beef breeds (*Manuscript in preparation*). Seven animals were sequenced with PacBio HiFi, five with Oxford Nanopore Technology (ONT), and 163 with PacBio CLR (sup. table 1). One sample, Charolais heifer, was sequenced using the three LR sequencing technologies, available in European Nucleotide Archive (ENA) [35].

To search for SVs, the LR data were first aligned against the ARS-UCD1.2 bovine reference genome [29] using the PacBio pbmm2 [48] with default parameters to generate a BAM file for each sample. Secondly, we searched for SVs using three different SV detection software, namely CUTESV v2.0.2 [36], PBSV v2.6.2 [37] and SNIFFLES v2.0.7 [38]. For both PBSV and SNIFFLES, we used default parameters. For CUTESV, we used the parameters *-- max_cluster_bias_INS 100 --diff_ratio_merging_INS 0.3 --max_cluster_bias_DEL 200 -- diff_ratio_merging_DEL 0.5* allowing insertions to be clustered within 100 bp and deletions within 200 bp during the SV call.

Third, we filtered the SV results in order to select only bi-allelic insertions and deletions between 50 nucleotides and 100 Kb in length, with a quality score greater than 20, well-defined SV breakpoints, and located on the 29 autosomes and the X chromosome.

### Benchmarking metrics

We used the TRUVARI benchmarking tool [39] to compare the base set, the putative true set of SVs, and the comp set, a set of SVs under evaluation. We applied the following parameters: a maximum distance between SVs of 500 bp (*–refdist 500*), a minimum size similarity between SVs of 0.7 (--*pctsize 0.7*), SV types considered during comparison (*-- typeignore FALSE*), no reciprocal overlap (*--pctovl 0*), and sequence similarity ignored (*-- pctseq 0*). This last parameter allows comparison of SVs generated by tools that do not explicitly report reference and alternative sequences in their VCF files. Our analyses are based on the summary output metrics from truvari, including: recall (the proportion of SVs in the base set that are recalled in the comp set: *TPbase*⁄(*TPbase + FN*)), precision (the proportion of SVs in the comp set that are found in the base set: *TPcomp⁄*(*TPcomp + FP*)), F1 score (combining the recall and precision: *2\** ((*recall * precision*)*⁄*(*recall + precision*))), and genotype concordance (the proportion of correctly genotyped SVs among all true positives, *TPcomp*_*TPgt*_ *⁄*(*TPcomp*_*TPgt*_ + *TPcomp*_*FPgt*_)).

### Evaluation of SV detected from different callers and LR data

We used BCFTOOLS v1.9 [44] to extract SV information, including length, calling support, genotype quality, and read depth, from CUTESV, PBSV and SNIFFLES based on the three LR data of the Charolais heifer. Here we used PacBio HiFi reads as the base set because of their high quality. We identified consensus SVs that were called by CUTESV, PBSV and SNIFFLES on HiFi data in two steps. First, we benchmarked the SVs called by PBSV and CUTESV to find the intersect SVs. Then, the intersect SVs were benchmarked against SVs called by SNIFFLES. To evaluate the performance of CUTESV, PBSV, and SNIFFLES across different LR technologies, we benchmarked SVs called on HiFi, as the base set, against SVs called on ONT or CLR, as the comp set. The benchmarking was performed separately for the three callers of CUTESV, PBSV, and SNIFFLES.

For the ensemble calling of the three tools, we applied intersect and union approaches. For the former, we used the TRUVARI benchmark to obtain *TP-base* as the ensemble SVs. While for the union, we used JASMINE v1.1.5 [41] with default parameters to merge the intended sets of SVs.

Following the developer’s recommendation to increase sensitivity and recall, we applied the *- -tandem-repeats* parameter in the SV discovery step of PBSV. We tested the performance on three HiFi samples from the Abondance (ABO), Tarentaise (TAR), and Vosgienne (VOS) breeds. As the input, we used a publicly available tandem repeat annotation [49] and our own generated annotation set. For the latter, tandem repeat regions were inferred using TRF v4.09.1 [50] on the bovine reference genome ARS-UCD1.2 [29]. We applied recommended parameters set to a match weight of 2, a mismatch penalty of 7, an indel penalty of 7, a match probability of 80, an indel probability of 10, a minimum alignment score of 50, and a maximum period size of 500.

### Evaluation of short reads SV genotyping tools

Illumina short reads (SR) of the Charolais heifer were available at 112x. SR data were also available for 154 samples (*Manuscript in preparation*). We down-sampled the coverage of the Charolais heifer using SEQTK [51] to 60, 30, 15, and 10X, respectively, to mimic the coverage of other samples. SR from this heifer and the other 154 samples, for which both LR and SR were available, were then processed subsequently to produce a BAM file for each sample. SR sequences were aligned against the ARS-UCD1.2 [29] reference genome with BWA MEM [52], duplications were removed and chromosomes were sorted using SAMTOOLS v1.19 [44], and base quality score recalibration was performed using GATK v4.5 [53] based on the known ∼106 million SNPs and InDels from the 1000 Bulls genome project. The BAM file was then processed by DELLY v1.2.6 [10], LUMPY [11] via SMOOVE v0.2.8 [54], and MANTA [9] to call SVs. We only retained deletions called by LUMPY and DELLY with the additional PASS flag for DELLY. We kept both deletions and insertions produced by MANTA.

For each sample, the SVs detected by PBSV on the CLR data were used as the base set. This base set was also used as known SV sites to genotype from the SR data using GRAPHTYPER v2.7.7 [21], PARAGRAPH v2.3 [22], SVTYPER v0.7.1 [23] and Variation Graph toolkit (VG) v1.56 [24]. The first three tools take a BAM file as input, the same as used by LUMPY, DELLY and MANTA. With default parameters, GRAPHTYPER generates numerous chunks of genotyped SVs across the genome. We used the BCFTOOLS *--naive* concat function to combine all the chunks, sort them and retain only biallelic SVs defined by the “*AGGREGATED*” SV model with a size greater than 50 bp and a maximum alternative allele support of 0.5. For PARAGRAPH, we used information on the average SR length and alignment coverage from the final BAM file. For SVTYPER, we added information from *CIPOS* and *CIEND* with values of -*100,100*; and *CIPOS95* and *CIPOS95 END* with *0,0* to the vcf info field of known SV sites. We genotyped SVs using svtyper-sso with the parameters *--max_reads 1000000 --max_ci_dist 4 -- split_weight 1 --disc_weight 1*. We then removed all SVs with missing genotypes defined by PARAGRAPH and SVTYPER.

For SV genotyping using VG in each sample, we first constructed the variation graph for all chromosomes separately based on the individual’s base set. These separate chromosome graphs were then merged into a whole-genome graph by re-setting their node ids to follow the order of the 29 autosomes plus the X chromosome. The whole-genome graph was then indexed to an *.XG*, converted to *.GFA* and libraries (*.GBZ*, *.MIN*, and *.DIST*) were generated to facilitate the alignment. SR were mapped against the graph using *VG GIRAFFE* [55] to create a GAM file. The alignment and graph were chunked by chromosome. *SNARLS*, which contains breakpoint information of SVs embedded in the graph, and *PACK,* which contains the post-alignment coverage index with a minimum mapping quality of 5 (*--min-mapq 5*), files were created from the chunked graph and alignment, respectively. SV genotyping was then performed based on the evidence in these *SNARLS* and *PACK* files. We retained only biallelic genotyped SVs across autosomes and X chromosome. By default, VG only outputs SVs with heterozygous or homozygous alternative genotypes. As the output does not include SVTYPE information, we classify genotyped SVs as deletions if the REF allele is longer than 50 bp and the ALT allele is one bp, or as insertions if the REF allele is one bp and the ALT allele is longer than 50 bp.

We performed benchmarking on the genotyped SVs recovered from SR by seven tools, using them as comp sets against the base set for each individual. The benchmarking was carried out separately for deletions and insertions.

### Extending the number and diversity of reference SV panel

We divided the 151 samples with available LR and SR into validation and reference sets. The validation samples consisted of two individuals each from Holstein (HOL), Montbéliarde (MON), and Normande (NMD). From the 145 samples in the reference set, we created the first reference panel by merging the base sets (SV discovered by PBSV on CLR data) of 23 HOL samples. The second panel was created by adding the base sets of 23 MON reference samples. The third panel was created by adding the base set of 24 NMD reference individuals. And the last panel consisted of all reference samples, resulting in a reference panel of 145 samples from 14 breeds. Base sets of validation samples were not included in the creation of any of these panels.

We used JASMINE v1.1.5 [41] to merge the base sets of reference samples with the default parameters *--ignore_strand --mutual_distance --allow_intrasample --output_genotypes* for each panel. For each panel, only SVs with a minimum number of alternative alleles of four were retained using VCFTOOLS [56]. The SRs of the validation samples were aligned to the graphs and the genotyped SVs were called. The genotyped SVs, here as comp sets, were then compared against the base set of each validation individual. We used the deconstruct option of VG for decomposing the graph referring to positions in the ARS-UCD1.2 and counted the bubbles generated within.

### Optimum parameters for building a SV reference panel of 14 breeds

We kept the 6 validation and 145 reference samples from the previous section. Apart from the default JASMINE parameters, we also modified the merging parameters with *-- max_dist_linear=0.1* and *--min_dist=50*. In short, compared to the default parameters, JASMINE only merged SVs within a distance proportional to 0.1 of their size, instead of 0.5, and with a minimum distance of 50, instead of 100. These parameters were recommended by the JASMINE developer in a github issue and increased the stringency of SV merging.

In addition to the minimum alternative-allele counts of four, we also applied eight thresholds of 2, 3, 5, 7, 13, 20, 27, and 36 from VARCALLS after SV merging. This information is output by JASMINE and indicates the number of variant calls supporting the merged SVs. Thus, 8 x 2 reference panels were created from these adjustments, on which variation graphs were constructed. Subsequently, the SR data of the validation samples were aligned to these variation graphs and subjected to VG genotyping. When applying VARCALLS thresholds, we removed SVs from the reference panel. Subsequently, we adjusted the base set of validation samples by also excluding the SVs that were removed due to VARCALLS filtering. We benchmarked the genotyped SVs (as comp sets) against the adjusted base set of validation samples using TRUVARI.

### Genotyping SV on a large cohort of individuals sequenced with short reads

We created the final reference panel using the SV sets called by PBSV on LR of 176 samples from 14 breeds (Table 1, sup. table 1). We merged the SVs with JASMINE using default parameters with a VARCALLS threshold of seven, followed by the creation of the final variation graph. SR from 571 individuals were aligned against the graph and then processed for SV genotyping. The collection and processing of these SR samples has been described elsewhere (*Manuscript in preparation*). We used the --*Aa* parameter in the VG call to output all genotypes, including missing or homozygous reference, for all SVs in the final panel. Annotation of the genotyped SVs was carried out using snpEff [42] based on the bovine annotation of ARS-UCD1.2 v99.

Principal component analysis (PCA) was performed on the genotyped SVs using PLINK v1.9 [57] with 20 eigenvectors. For 148 samples with LR and SR available, we benchmarked the genotyped SVs, as the comp set, against the base set of each sample, respectively. We used VCFTOOLS [56] to infer the frequency of genotyped SVs. We used UpSet [58] to characterize major (AF>0.7) or rare (AF<0.1) SVs in all samples. We used BEDTOOLS [59] to intersect SVs regions of interest to repeat annotations [49] of the bovine reference genome. R [60] and GGPLOT2 [61] were used mainly for data handling and plotting. The pipelines used in the analysis have also been made publicly available (see *Data availability*).

## DATA AVAILABILITY

The <u>S</u>nakemake script used in the analysis is available at https://github.com/mas-agis/cascad1/. All the raw data will be made available once manuscript is accepted for publication.

## SUPPLEMENTARY DATA

Additional file 1. Supp. Figure 1-15.

Additional file 2. Supp. Table 1-11.

## AUTHOR CONTRIBUTIONS

MMN, TF, CK, DB, MPS, and MB conceived the study. MMN performed the analysis and wrote the original draft. TF, CK, DB, MPS, MB contributed to writing the manuscript. CeG and SF collected samples and realized extraction. CE, CM, AS, CD, ChG, C.I. CK, CV, and DM conceived the experimental design and supervised the technical aspects of the project and data deposition in public databases. ADF, CB, and TF conducted SV detection. All authors reviewed the final manuscript.

## ACKNOWLEDGEMENTS

We are grateful to the genotoul bioinformatics platform Toulouse Occitanie (Bioinfo Genotoul, https://doi.org/10.15454/1.5572369328961167E12).

We also thank the INRAE GeT-PlaGe facility for whole genome sequencing. GeT-PlaGe is a member of France Génomique national infrastructure, funded as part of the “Investissements d’Avenir” program managed by the Agence Nationale de la Recherche (contract ANR-10-INBS-09).

