## Supplementary material for "Comprehensive detection of structural variations in long and short reads dataset of French cattle": Suplemental Figure 1-15: draft_clean_supFig.docx

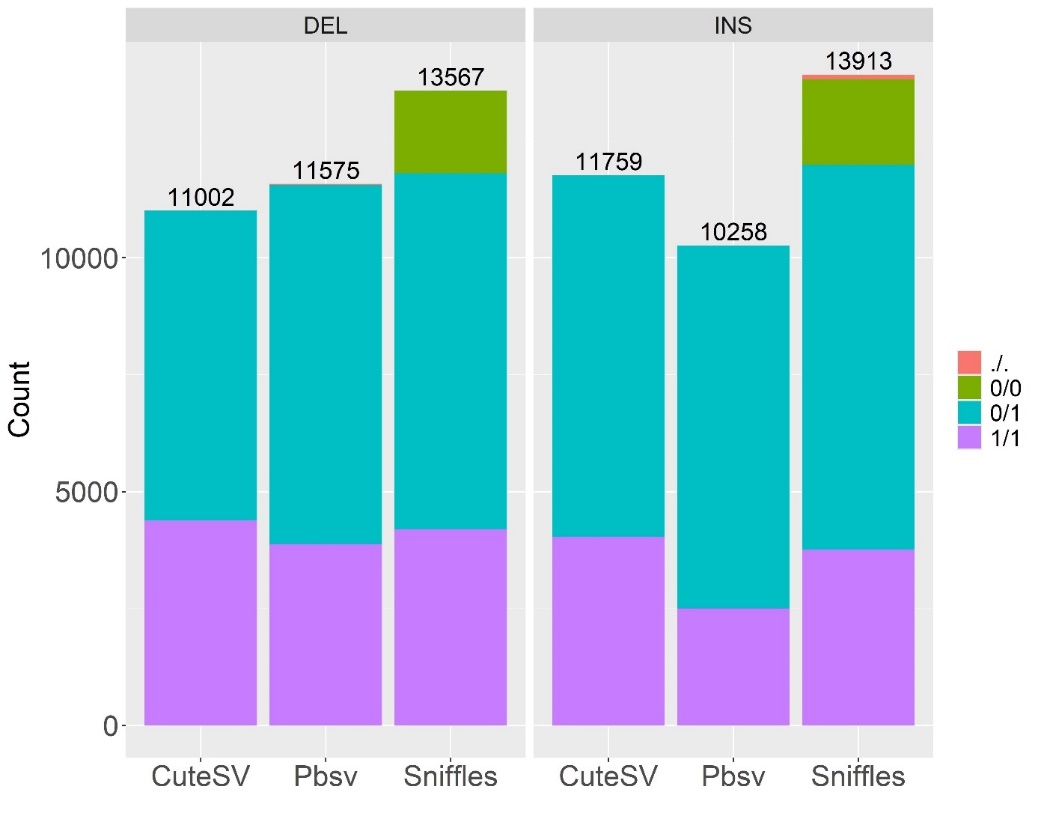

Supplemental figure 1. Genotype distribution of SV detected on HiFi data using Cutesv, Pbsv, and Sniffles.

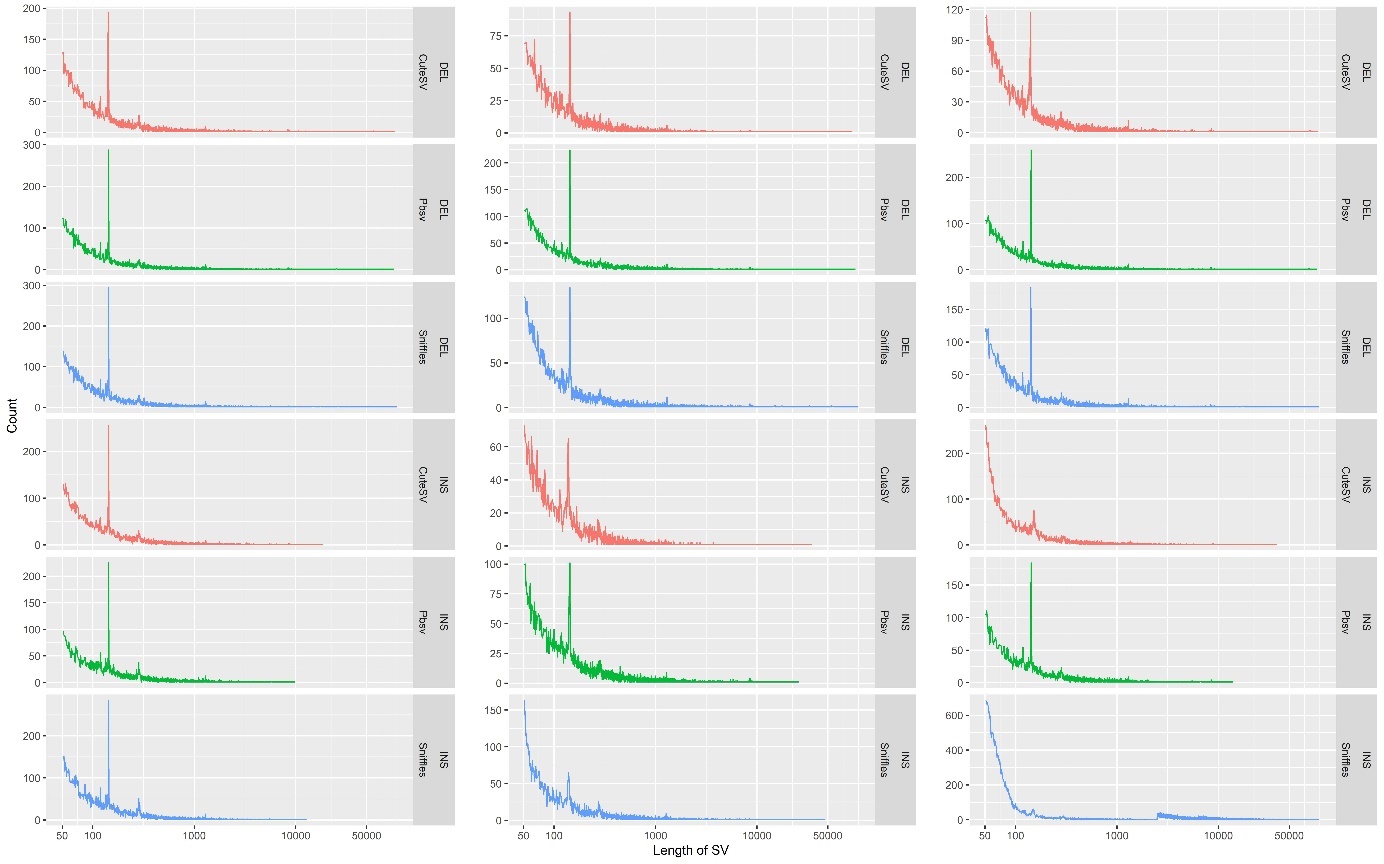

Supplemental figure 2. Distribution of SV length across genome of the Charolais heifer (re-creation of main figure 1b separately for each caller, svtype and long-reads data). First column is for Pacbio-HiFi, second for Oxford-ONT, and third PacBio-CLR.

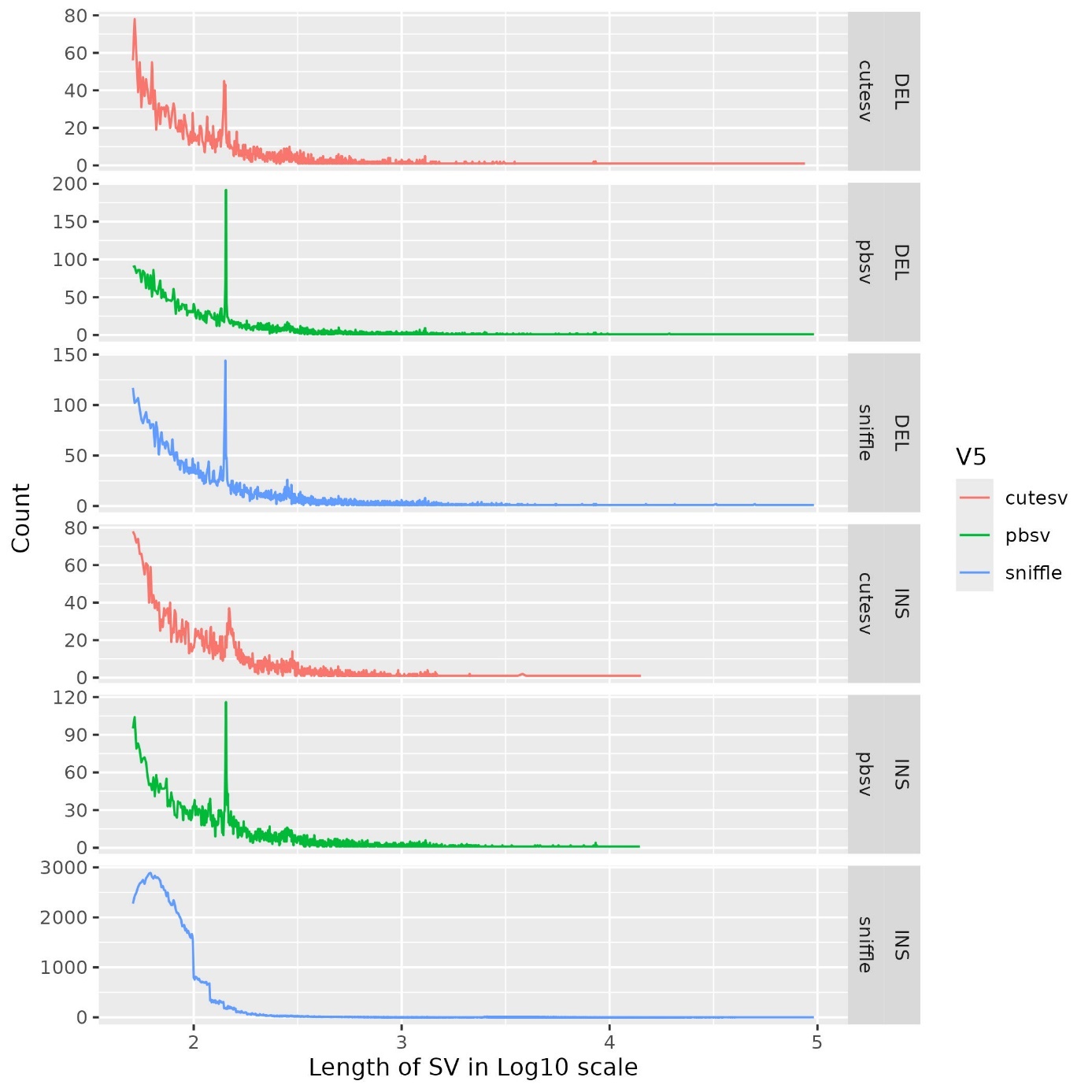

Supplemental figure 3. Distribution of SV length across genome of HOL28 sequenced with PacBio CLR. The first three rows are for deletions and the last three rows for insertions. The legends indicate the tools used in discovery of SVs (CuteSV, Pbsv, and Sniffles).

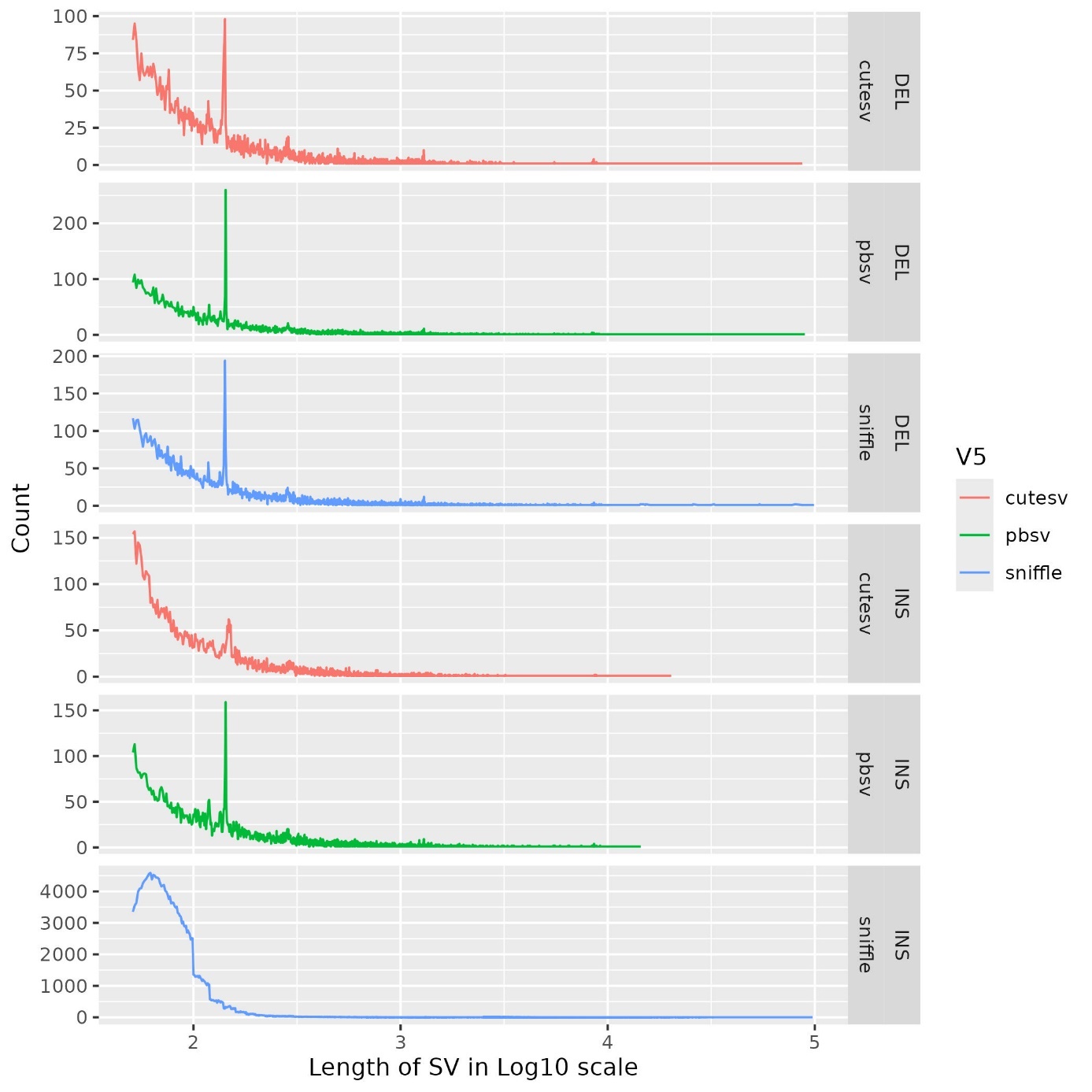

Supplemental figure 4. Distribution of SV length across genome of MON16 sequenced with PacBio CLR. The first three rows are for deletions and the last three rows for insertions. The legends indicate the tools used in discovery of SVs (CuteSV, Pbsv, and Sniffles).

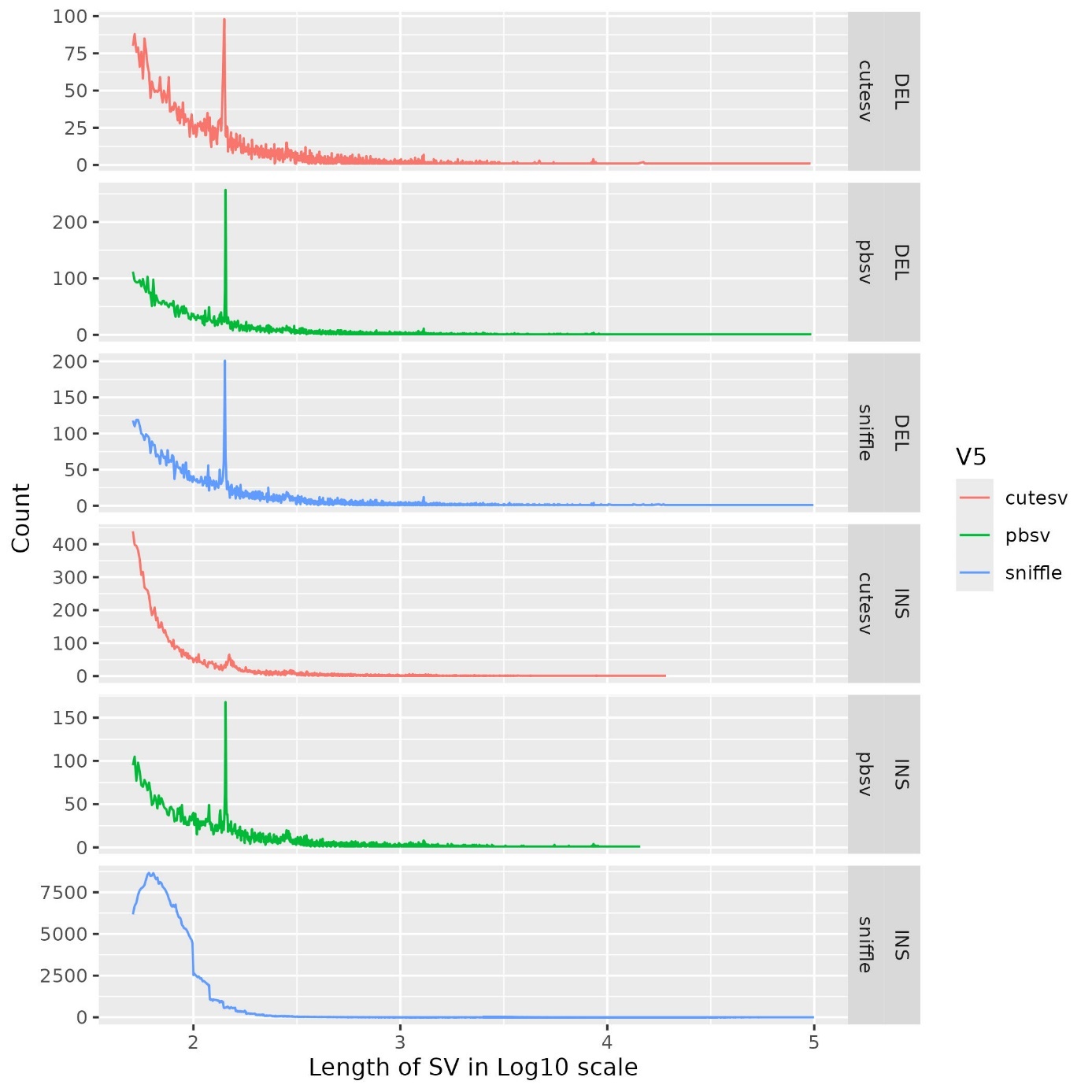

Supplemental figure 5. Distribution of SV length across genome of NMD19 sequenced with PacBio CLR. The first three rows are for deletions and the last three rows for insertions. The legends indicate the tools used in discovery of SVs (CuteSV, Pbsv, and Sniffles).

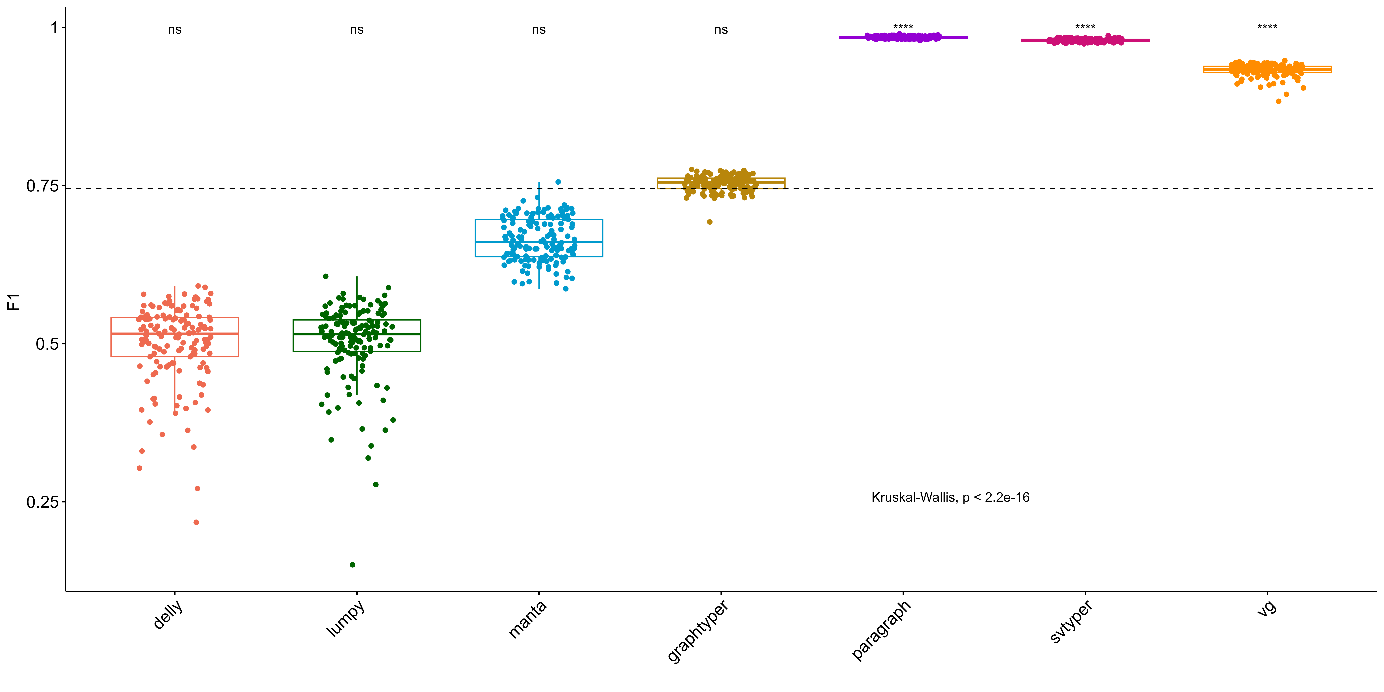

Supplemental figure 6. One-sided Kruskall-walis test on the F1 score outputted by seven SV genotyping tools on deletion of 148 samples. The dashed line indicates the average F1 score across all genotyping tools.

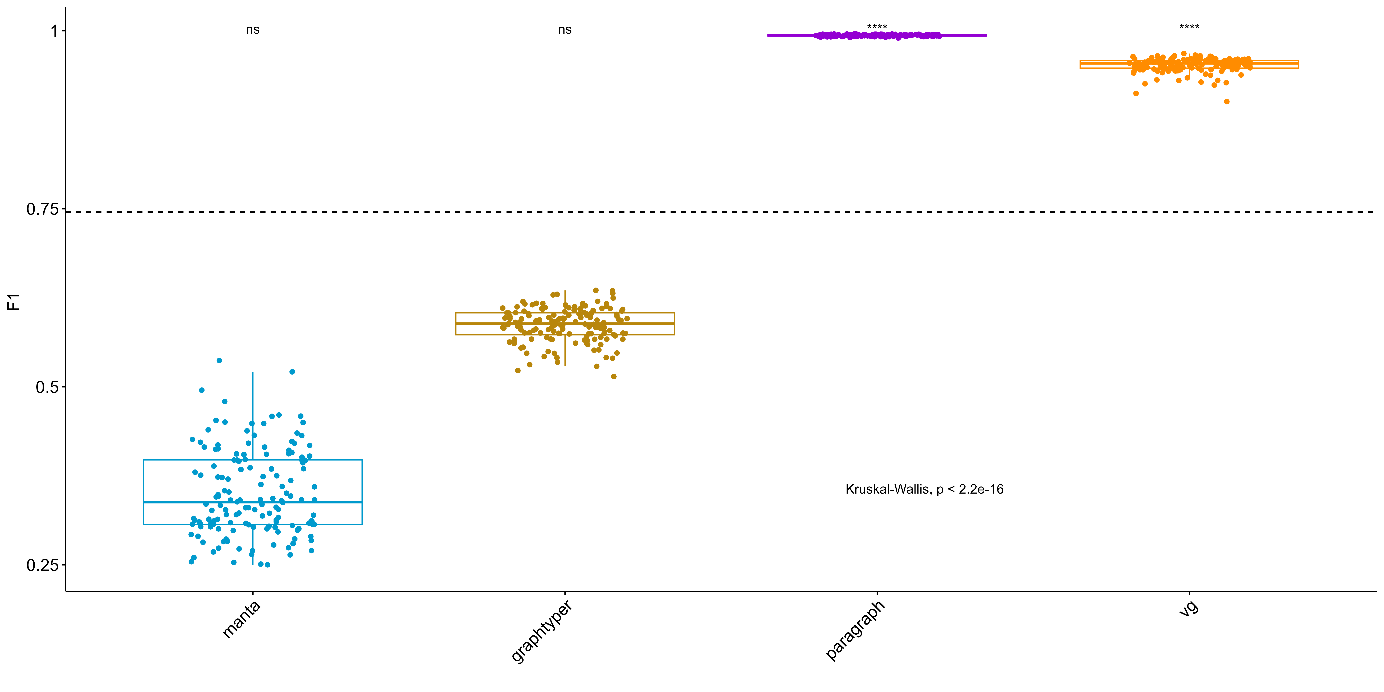

Supplemental figure 7. One-sided Kruskall-walis test on the F1 score outputted by four SV genotyping tools on insertion of 148 samples. The dashed line indicates the average F1 score across all genotyping tools.

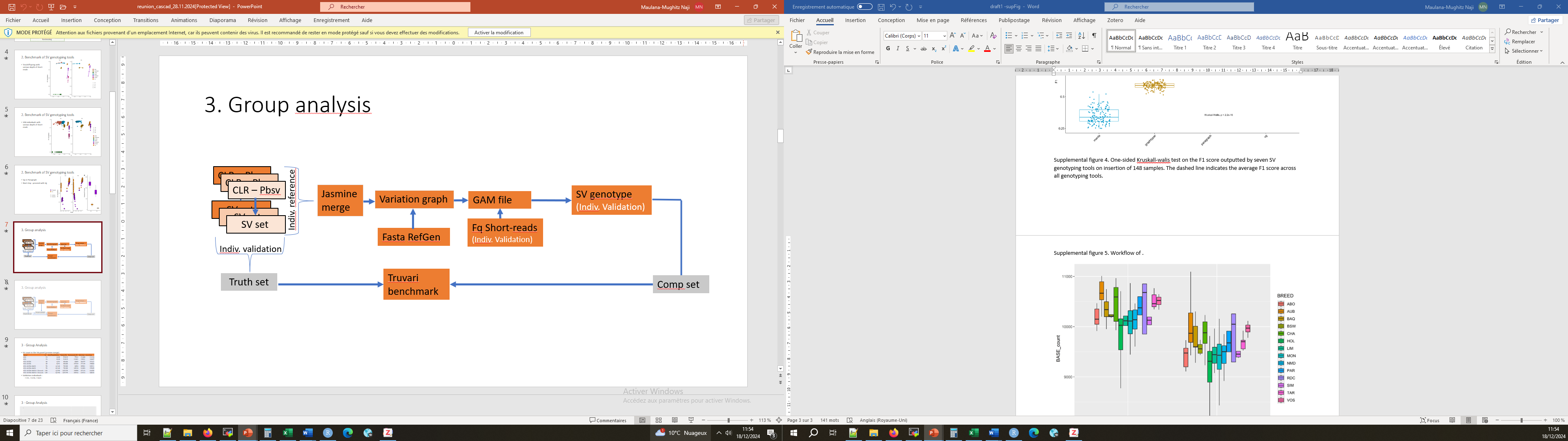

Supplemental figure 8. Workflow of genotyping SV from short-reads on various SV reference panels.

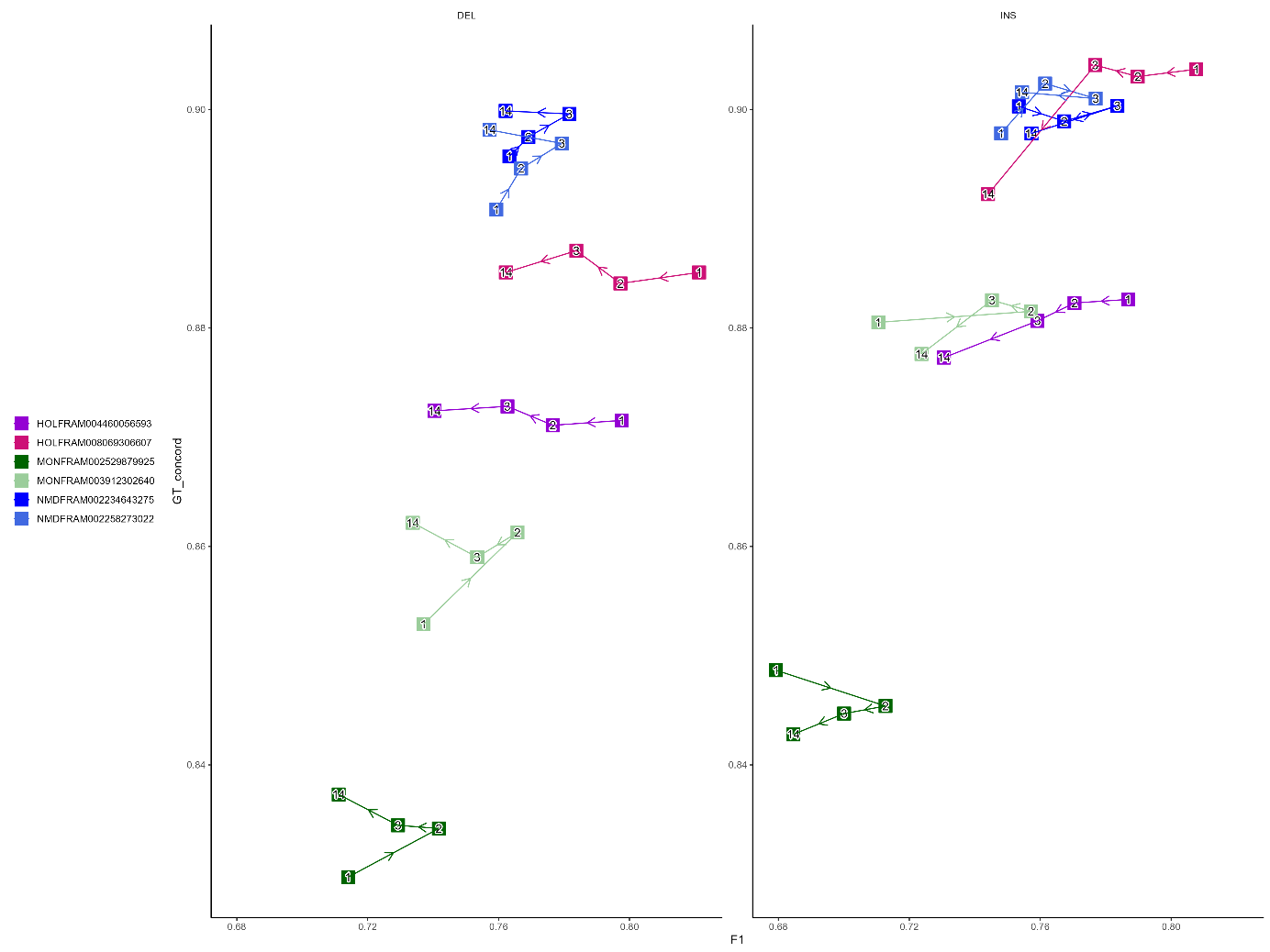

Supplemental figure 9. F1 score and genotyping concordances of genotyping structural variants (SV) of 6 validation individuals using short-reads respectively on variation graphs created from different panels of individuals and breeds.

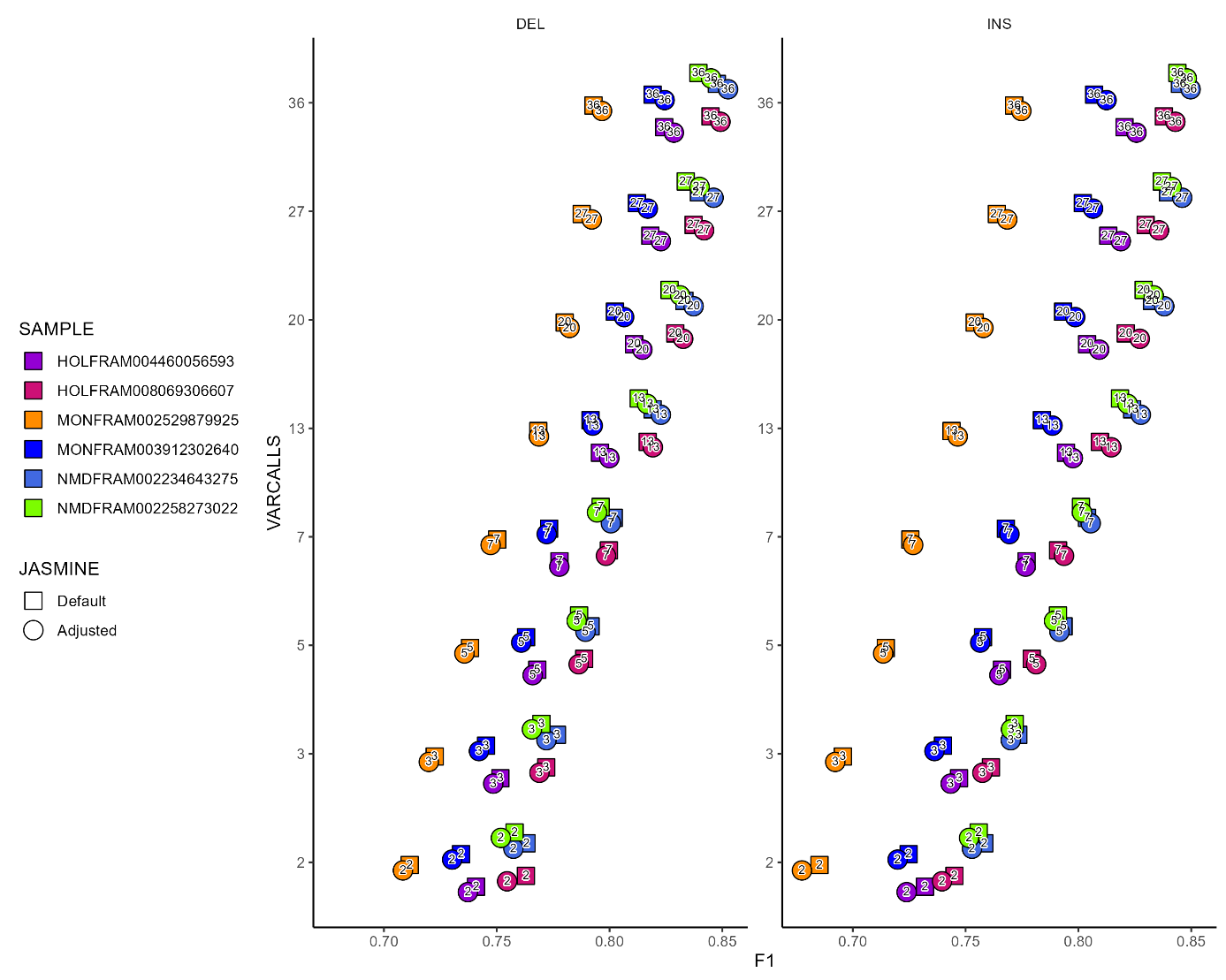

Supplemental figure 10. F1 score of genotyping structural variants (SV) of 6 validation individuals on variation graphs created from 145 samples of 14 breeds by default and adjusted jasmine merging parameters combined with different VARCALLS thresholds of 2, 3, 5, 7, 13, 20, 27, 36 as indicated in the graph points.

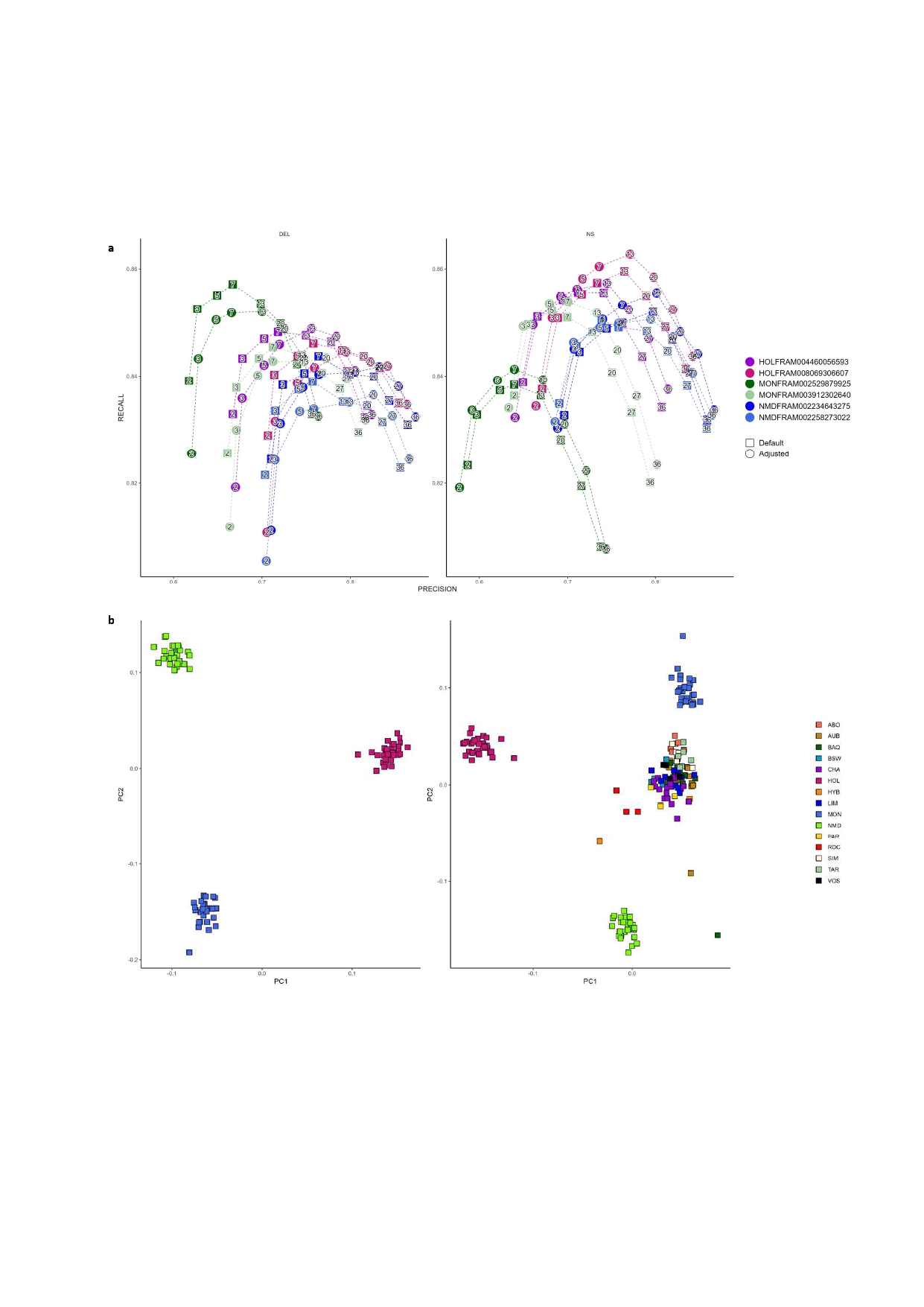

Supplemental figure 11. Precision and recall rates of genotyping structural variants (SV) of 6 validation individuals on variation graphs created from 145 samples of 14 breeds by default and adjusted jasmine merging parameters combined with different VARCALLS thresholds of 2, 3, 5, 7, 13, 20, 27, 36 as indicated in the graph points.

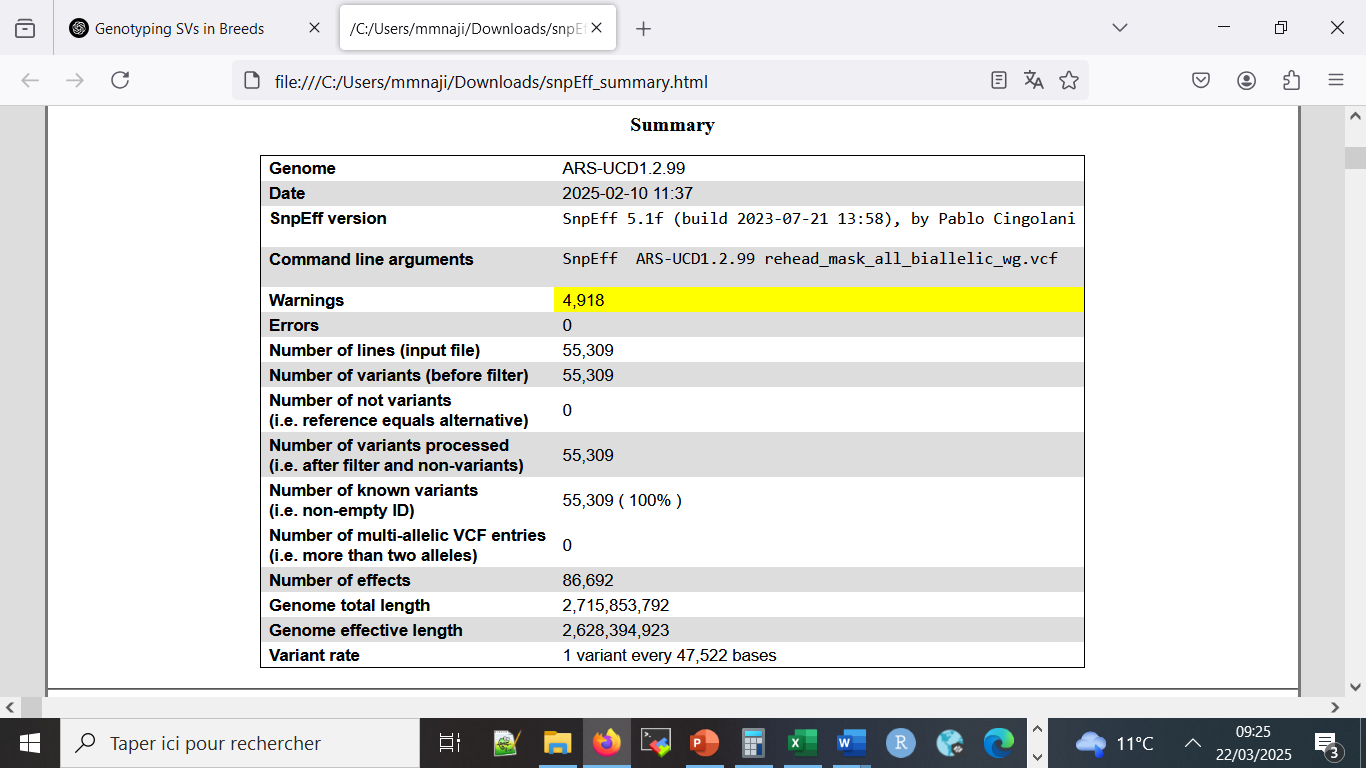

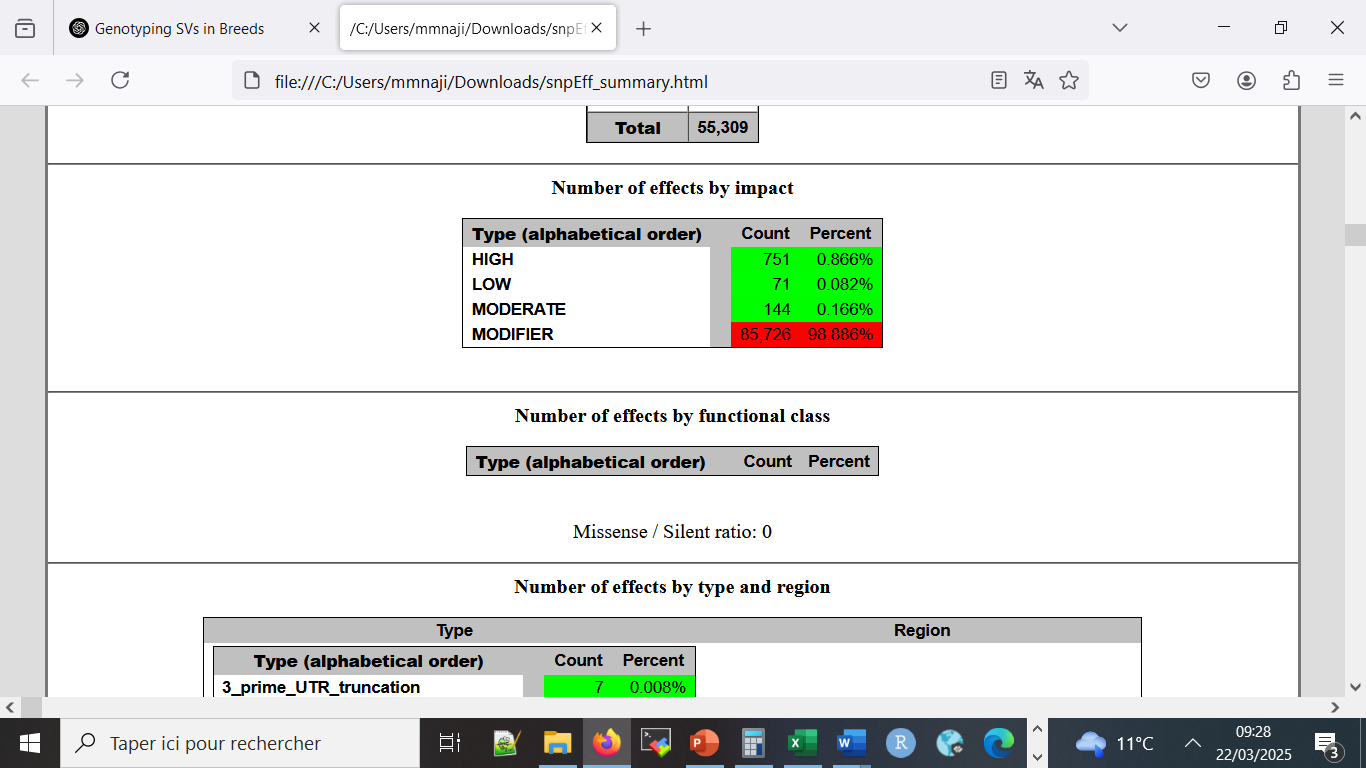

Supplemental figure 12. Annotation of genotyped SVs from the final reference panel. Left: Annotation summary. Right: Number of annotated effects by impacts.

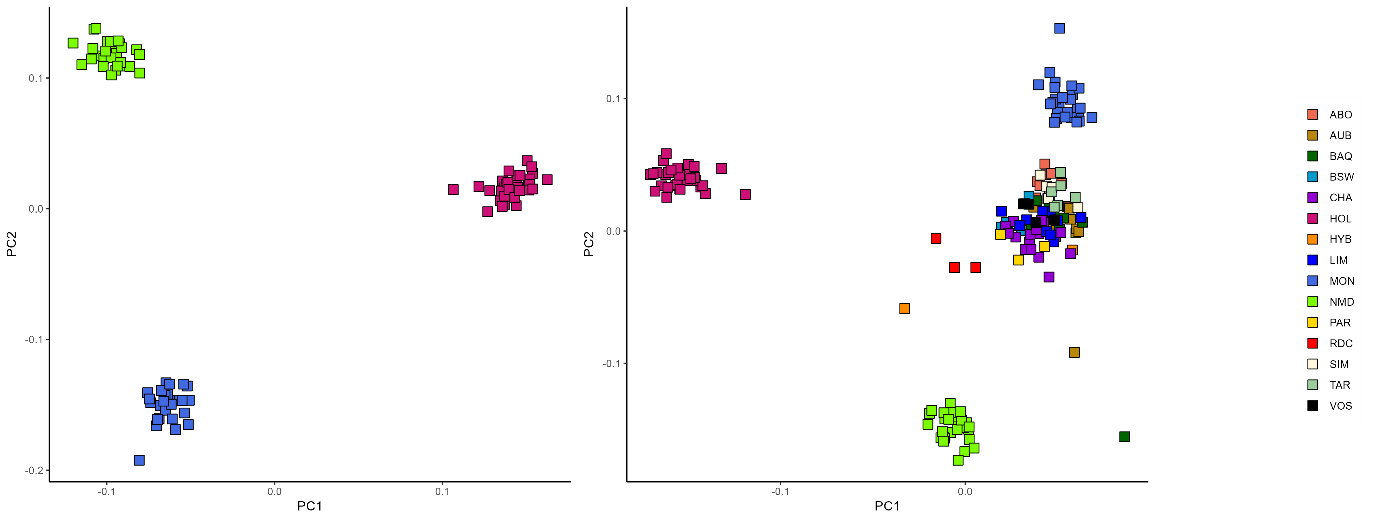

Supplemental figure 13. Principal component analysis (PCA) of the final SV reference panel, constructed from 176 long-read samples using the default Jasmine merging parameters and a VARCALLS threshold of seven. **Left:** PCA performed on three breeds—Holstein, Montbéliarde, and Normande—with 147, 91, and 64 long-read sequenced samples, respectively. **Right:** PCA conducted using all 14 breeds with long-read sequencing.

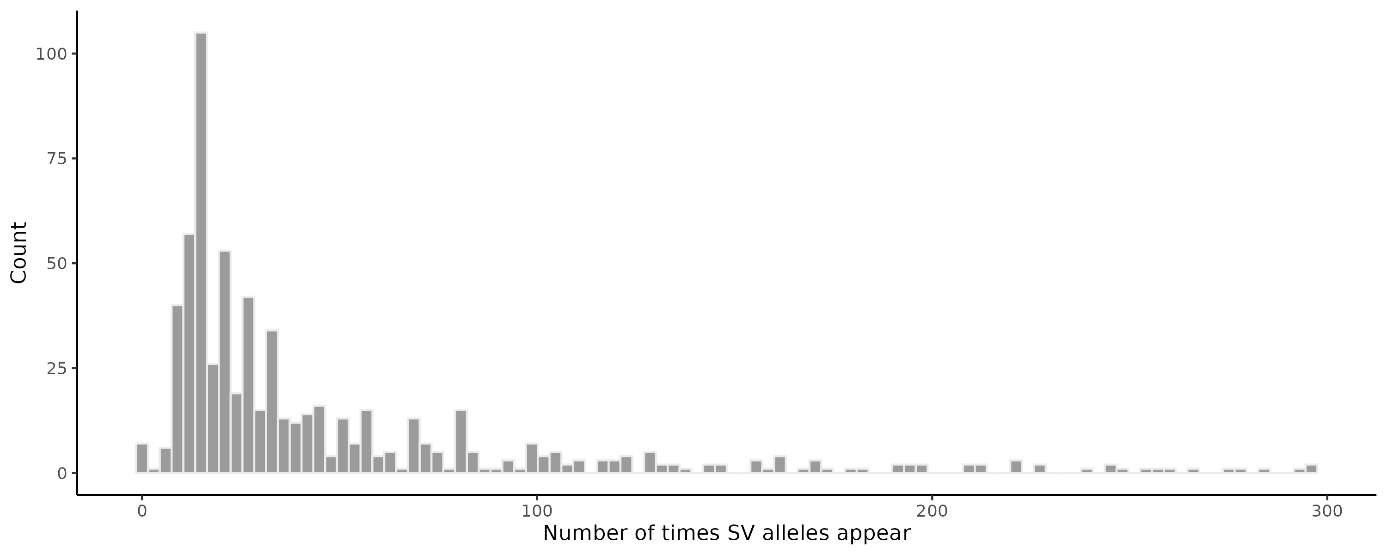
 Supplemental figure 14. Histogram of homozygous-reference SVs, genotyped across 571 short-read samples, that were initially called in the base set of 148 samples with both LR and SR available.

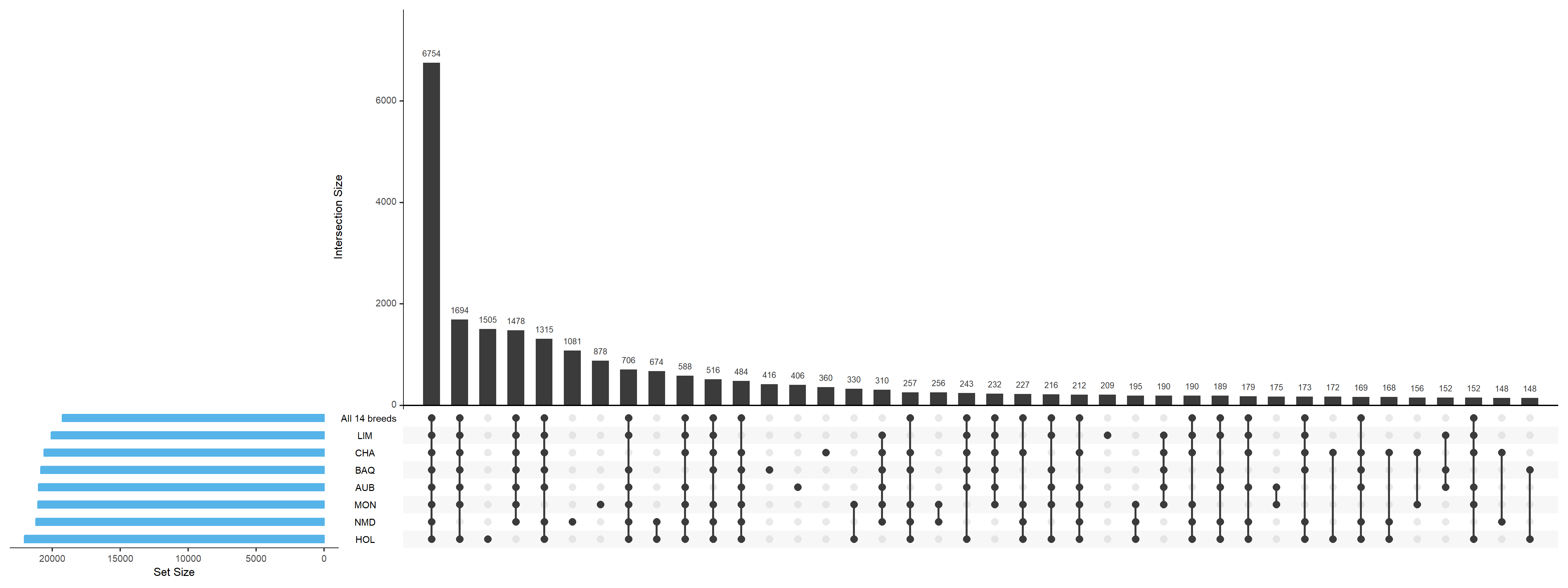

Supplemental figure 15. Upset plot of rare SVs (allele frequency < 10%) across all 14 breeds and six breeds with short-read samples higher than 30.
